# Nonapeptide molecular evolution during the adaptive radiation of Tanganyika cichlids

**DOI:** 10.1101/2025.06.18.660417

**Authors:** Pol Sorigue, Walter Salzburger, Rui F. Oliveira

## Abstract

Oxytocin (OT) and vasotocin (VT) are evolutionarily conserved nonapeptides that regulate a wide range of physiological and behavioural processes in vertebrates. Their receptor families have undergone gene duplications that facilitated functional diversification throughout vertebrate evolution. Using the diverse cichlid species in Lake Tanganyika, which have undergone repeated evolutionary transitions between social phenotypes, we investigated the molecular evolution of the nonapeptide system and its potential involvement in social behaviour. We performed a positive selection analysis based on the dN/dS ratio and examined the correlation between amino acid variants and two social phenotypes. We also analysed gene expression data to explore associations between brain receptor expression and social phenotype variation. Our findings reveal that, while most sites in nonapeptide receptors are under strong purifying selection, a few sites - primarily in the extended intracellular loop 3 (IL3) of VTR2A receptors - show signatures of positive selection. Additionally, a specific amino acid in VTR2Aa correlates with pair-bonding, suggesting its potential role in social attachment. Together, our results provide new insights into the evolution of the nonapeptide system and its contribution to social diversity in cichlids.

## Introduction

Oxytocin (OT) and vasotocin (VT) are two closely related nonapeptides that play essential roles in various physiological processes across vertebrates. Derived from a common ancestral molecule, these peptides share a highly conserved structure and function alongside two evolving families of receptors. Gene duplications have expanded receptor diversity, enabling the acquisition of new functions over evolutionary time. OT and VT have been extensively studied for their roles in osmoregulation, reproduction, and social behaviour (Balment et al., 2006; Jurek & Neumann, 2018), yet their precise mechanisms of action and evolutionary adaptations remain incompletely understood.

An ancestral VT-like peptide and its cognate receptor existed before the emergence of vertebrates. This ancient VT-like peptide played key roles in the regulation of reproduction and in diuretic functions (Kobayashi et al., 2022; Oumi et al., 1994). Around 540 million years ago (mya), the ancestral gnathostome vasotocin receptor (*VTR*) gene underwent a local duplication giving rise to a *VTR1*/*OT* receptor (*OTR*) and a *VTR2* (Gwee et al., 2009; Sartorius et al., 2024). In turn, a segmental duplication of *VT* gave rise to *OT* around 530 mya (Banerjee et al., 2017; Sartorius et al., 2024)(Banerjee et al., 2017; Sartorius et al., 2024), leading to a contiguous head-to-tail genomic arrangement of the two ligand genes. This disposition is common across vertebrates, with the exception of mammals, in which the orientation is tail-to-tail so that the genes are transcribed from different strands; also, in some teleosts, the two nonapeptides are found on separate chromosomes (Gwee et al., 2009). The receptor evolution in vertebrates was determined by two rounds of whole genome duplication (1R, 2R) followed by clade-specific losses (Ocampo Daza et al., 2022; Theofanopoulou et al., 2021). The teleost-specific third whole genome duplication (3R) event expanded the receptor repertoire in this group, which now includes two OTRs and up to seven VTRs, classified into four receptor types: VTR1A, VTR2A, VTR2B and VTR2C (Mayasich & Clarke, 2016; Ocampo Daza et al., 2022; Theofanopoulou et al., 2021).

Nonapeptides are encoded by precursor genes that usually span about 2,000 base pairs. The precursor genes typically contain three protein-coding exons, and they produce proteins that undergo modifications to yield the final nonapeptide hormone. The protein sequence is over 100 amino acids long, and comprises a signal peptide, hormone moiety, and neurophysin domain, with an additional copeptin domain in vertebrate VT and teleost OT (Ivell & Richter, 1984). The cyclic structure of nonapeptides is conserved across vertebrates, with variation mainly at amino acid positions 3, 4, and 8. Their classification into two families—basic nonapeptides (VT family) and neutral nonapeptides (OT family)—is based on the nature of the amino acid at position 8. Despite phylum-specific sequence variation, several residues within the nonapeptide sequence remain highly conserved (Banerjee et al., 2017). In teleosts, the amino acid sequences of VT and OT nonapeptides are homologous, except at positions 4 and 8 (Gimpl & Fahrenholz, 2001). Neurophysin, a crucial carrier protein, ensures peptide stability, folding, processing and storage prior to nonapeptide release into the bloodstream (Grigor’eva & Golubeva, 2010). It contains 14 highly conserved cysteine residues essential for the proper folding of the precursor protein, and is linked to the hormone via a conserved glycine-lysine-arginine (GKR) sequence. Finally, the copeptin domain, present in vertebrate VT and teleost OT, has an unclear function but may be involved in prolactin release (Flores et al., 2007).

OTRs and VTRs are G protein coupled receptors (GPCRs) with seven transmembrane α-helices domains (TM1-7), alternated by three intracellular loops (IL1-3) and three extracellular loops (EL1-3). The transmembrane domains form a ring structure that causes the extracellular domains to confer a cavity where the nonapeptide binds (Mouillac et al., 1995; Waltenspühl et al., 2022). The N-terminal domain is exposed to the extracellular space and is also involved in ligand binding, while the intracellular C-terminus interacts with the G protein for intracellular signal transduction (Barberis et al., 1998). Generally, OTRs and VTRs share high sequence similarity. General features, as well as some functionally crucial amino acid residues, are conserved across distant lineages and receptor types (Gimpl & Fahrenholz, 2001). Most nonapeptide receptors in vertebrates are encoded by two exons. However, the teleost ancestral *VTR2A* gene accumulated three more introns that led to the common 5-4 exon-intron organization of the two teleost *VTR2A* gene copies *VTR2Aa* and *VTR2Ab* (Ocampo Daza et al., 2022). Furthermore, teleost *VTR2A* copies have an extended intracellular loop 3 that contains several residues that can be phosphorylated, making it a potential substrate for functional divergence (Ocampo Daza et al., 2022). OTRs, VTR1s, VTR2Bs and VTR2C couple mainly with G_Q_ activating the calcium-mobilizing mechanism as a second messenger for downstream signalling through phospholipase C pathway (Brinton et al., 1994; Yamaguchi et al., 2012). In contrast, VTR2A couples with G_S_ and activates cAMP pathway (Wang et al., 2021; Zhang et al., 1997). Given that both receptors interact with a different region of the G protein, the structural difference might respond to distinct binding mechanisms (Waltenspühl et al., 2022). Mutagenesis analysis showed that a correct activation of G_Q_ requires a functional intracellular loop 2, while the intracellular loop 3 is necessary and sufficient for activation of G_S_ (Liu & Wess, 1996). Therefore, IL3 is a key player for VTR2A signalling, as it is essential for G_S_ activation and subsequent cAMP pathway activation.

Multiple gene duplications of the nonapeptide receptors facilitated the acquisition of new functions in vertebrates. Once duplicated, a gene can accumulate mutations without compromising the function of the original copy. This allows for the evolution of novel ligand specificities, signaling pathways, or tissue-specific expression patterns, leading to functional diversification (Glasauer & Neuhauss, 2014). Beyond its ancestral roles in reproduction and osmoregulation, the nonapeptide system acquired novel roles in vertebrates (Balment et al., 2006). This has been extensively studied in mammals (Jurek & Neumann, 2018), and growing evidence is showing that the nonapeptide system regulates multiple behaviours in teleost (Backström & Winberg, 2009; Landin et al., 2020; Ribeiro et al., 2020).

In teleosts, nonapeptide-secreting neurons have been found in the preoptic area (POA). Three subpopulations of OT and VT neurons – giganto-, magno-, and parvocellular neurons – project their fibers into diverse regions of the brain, as well as the spinal cord. The local expression of receptors largely determines the localised activity of nonapeptides across brain regions, but also determines functional connectivity across regions. Thus, the spatial and temporal distribution of receptors is crucial for modulating behaviour. Notably, OTRs and VTR1As have been found in all the brain areas of the teleost social decision-making network, an evolutionary conserved network, consisting of the mesolimbic reward system, the social behaviour network and shared areas connecting the two, that together regulate social behaviour (Goodson, 2005; Huffman et al., 2012; Kareklas et al., 2025; O’Connell & Hofmann, 2011). Although VTR2s have also been reported in the brain (Lema et al., 2019), their central functions are much less characterized. The nonapeptide system has been linked to various social behaviors in teleosts, including aggression, affiliative behaviour, parental care and pair-bonding (Godwin & Thompson, 2012; Nowicki et al., 2020; Reddon et al., 2017; Ripley & Foran, 2010). However, species-specific patterns make it difficult to establish clear roles for each nonapeptide and receptor. Instead, they function within a complex behavioural network influenced by multiple interacting factors, including expression levels, brain region, environmental context, and physiological state. Furthermore, growing evidence suggests that both OT and VT can bind to each other’s canonical receptors, influencing each other’s functions (Song & Albers, 2018). Therefore, despite a big effort has been done to assign specific behaviours to specific nonapeptides, their roles are often intertwined.

Cichlids are freshwater fishes known for their remarkable adaptive radiations, which have led to extensive speciation events (Ronco et al., 2021; Salzburger, 2018). The adaptive radiation of cichlid fishes in Lake Tanganyika comprises 243 endemic species, along with several riverine species (Ronco et al., 2020). These species are classified into 12 tribes and exhibit extensive phenotypic diversity, including variations in social behaviour. Multiple independent evolutionary transitions between social phenotypes occurred, which makes this system ideal for comparative studies. A variety of mating systems have evolved within the Lake Tanganyika cichlid radiation, which can be broadly categorized as monogamous and polygamous, and these are tightly linked to caregiving strategies. The emergence of monogamy has been proposed as a response to the need of biparental care of the offspring (Kleiman, 1977), although other studies see the latter as a consequence of the former (Stanbrook et al., 2022). The involvement of the nonapeptide system regulating social pair-bonding and parental care has been well documented in cichlids (O’Connell et al., 2012; Reddon et al., 2017). The present study aims to find molecular signatures in the nonapeptide system linked to the evolution of two social phenotypes across Lake Tanganyika cichlids. Whole-genome sequences are available for all species within the radiation, and transcriptomes exist for 74 species (El Taher et al., 2021; Ronco et al., 2021). In the present study, we first characterized the nonapeptide system across the Lake Tanganyika cichlid radiation. Then, we searched for signatures of positive selection potentially linked to social phenotypes and analysed the tissue-specific expression of nonapeptide-related genes to explore correlations between gene expression and social behaviour.

## Methods

An extended version of the Methods section can be found in Supplementary Methods.

### 1. Lake Tanganyika cichlids *de novo* assemblies

We used Illumina whole-genome raw reads from all cichlid species in Lake Tanganyika (Ronco et al., 2021) to create de novo assemblies for all available species. Genomic raw reads (Bioproject PRJNA550295; NCBI BioProject database) were mapped to the reference genome (Nile tilapia, *Oreochromis niloticus*; RefSeq accession GCF_001858045.2, female) using Bwa-mem (version 0.7.18; Li & Durbin, 2009) with default parameters. In Bioproject PRJNA550295, genomes for all the species of the radiation are available, ranging from 1 to 4 individuals per species. Transcriptomic raw reads (Bioproject PRJNA552202(El Taher et al., 2021) were mapped to the reference genome using STAR aligner (version 2.7.10a; Dobin et al., 2013). All six tissues available (brain, gills, lower pharyngeal jaw, liver, ovaries and testis) and 4 to 6 individuals across 73 species were used for the analysis.

### 2. Nonapeptide system genes repertoire

To identify the full repertoire of nonapeptide precursors and receptor genes in the reference genome, we used zebrafish (*Danio rerio*) and medaka (*Oryzias latipes*) genes. We obtained the protein sequences of the nonapeptides (Banerjee et al., 2017) and their receptors (Ocampo Daza et al., 2022), and we searched similar protein sequences in the reference genome using protein BLAST (“BLASTp”, https://blast.ncbi.nlm.nih.gov). All Nile tilapia’s sequences that matched the search (e-value < 1e-50) were kept for further analysis (Supplementary Table 1). To assign receptor types to Nile tilapia’s sequences we built a phylogenetic tree using IQTREE (version 2.0.3; (Nguyen et al., 2015) with the receptor sequences of distant teleost species (Supplementary Data 1). We assigned receptor type by phylogenetic proximity.

We investigated the presence and number of copies of all nonapeptide precursor and receptor genes in Lake Tanganyika cichlids. To this end, we used long-read PacBio assemblies (of six cichlid species representing six tribes: *Neolamprologus multifasciatus* (Lamprologini; GenBank assembly GCA_963576455.2, *Simochromis diagramma* (Tropheini; GenBank assembly GCA_900408965.1), *Bathybates minor* (Bathybatini), *Cyphotilapia frontosa* (Cyphotilapiini), *Cunningtonia longiventralis* (Ectodini), and *Cyprichromis leptosoma* (Cyprichromini) (unpublished). We created a local database with each PacBio assembly using ‘makeblastdb’ (version 2.16.0+). We performed nucleotide BLAST (“BLASTn”) searches with the reference gene sequences as query. In all six PacBio assemblies we found a clear match for each gene spanning the entire reference gene (e-value < 1e-200). The location of the nonapeptide system repertoire in the six PacBio assemblies can be consulted in Supplementary Table 2.

Because dN/dS positive selection analyses are based on protein-coding sequences, we searched for protein-coding sequences in the reference genes in the Nile tilapia RefSeq annotation. We analysed all the annotated splicing forms of all 10 genes, and downloaded the coding sequences of all the isoforms that differ to any extent in their protein sequence. To obtain the complete set of splicing forms available in Lake Tanganyika cichlids in these genes, we performed a protein BLAST search in NCBI database with the term “African cichlids” with all the nonapeptide system genes. This revealed an additional transcript of *OTRb*. A total of 16 splicing forms across the 10 genes were kept for further analysis. The coverage of all transcripts was calculated using Samtools depth (Danecek et al., 2021). Combining manual inspection of the BLASTn results obtained from the PacBio assemblies and the Illumina assemblies coverage files, we identified a 16 bp deletion in the fourth exon of VTR2Ab gene in multiple Lamprologini species. In addition, we identified four more genetic features that required further investigation (explained in Supplementary Methods). The consensus sequence of each gene was obtained for each individual using the genomic location of the coding sequences and the mapping files (BAM). We obtained a species consensus by merging the individual genes of the same species with Seqtk (https://github.com/lh3/seqtk/). To obtain the consensus from the BAM file, we used a combination of SAMtools and BCFtools (Danecek et al., 2021).

To assign each amino acid to a domain of the protein, we used InterPro (https://www.ebi.ac.uk/interpro/search/sequence/), a comprehensive database for predicting protein domains based on amino acid sequence. Using this tool, we obtained information on amino acid positions and domain architecture location along the peptide sequence.

### 3. Multiple Sequence Alignments

We built a multiple sequence alignment with MAFFT (version v7.526; Katoh & Standley, 2013) for each gene with the consensus sequences of all Lake Tanganyika cichlids and the reference gene from Nile tilapia genome. We used genomic and transcriptomic assemblies to improve the accuracy of the consensus. We then translated every coding sequence into amino acids, and checked for the presence of START/STOP codons, early STOP codons or sequence lengths not dividable by 3. We spotted multiple STOP codons with the sequence ‘TGA’. Although the codon ‘TGA’ is known as a translation STOP sequence, this also encodes a selenocysteine and does not necessarily stops the translation (Rajput et al., 2019). *VT* and *VTR2Bb.tr1* did not have a canonical START and we intentionally trimmed the alignment until the first methionine. Few additional manual changes were applied (discussed in Supplementary Methods). After the manual modifications, a codon-based alignment was then performed using PAL2NAL (version V14; Suyama et al., 2006) ensuring that positive selection analysis was computed on the correct reading frame.

We calculated the nucleotide diversity index (π) of the alignments using the Ape package (version 5.8) in R (version 4.4.0). To examine domain-specific variation, we calculated π for all domains only in transcripts that encode a canonical GPCR. We first calculated π for each of the individual domains and then we computed π grouping domains by type. Additionally, in order to spot possible evolutionary constraints in the genes, we calculated the rate of synonymous substitutions for all transcripts and all species using Yang and Nielsen method (Yang & Nielsen, 2000).

### 4. Positive selection

For each transcript, we built a ML phylogenetic tree using IQTREE (version 2.0.3; Nguyen et al., 2015). Among different substitution models, IQTREE automatically selects the best fit and creates a phylogenetic tree. We built gene trees with nucleotide and amino acid sequences. Whole-gene ω (dN/dS) values were obtained using HyPhy (version 2.5.62) with the Fixed Effects Likelihood (FEL) method (Kosakovsky Pond & Frost, 2005). We also used FEL to investigate the site-specific positive selection across the gene sequences of the nonapeptide system genes. Because the multiple sequence alignments showed variable dS values (Supplementary Data 3), we selected FEL as an appropriate method due to its ability to account for dS variation across sites and branches (Pond & Muse, 2005). We accepted a p-value of < 0.05 as a threshold of significance. We performed positive selection using gene trees and species trees for all transcripts.

### 5. Correlated evolution between discrete traits

We gathered phenotypic data for all species for which information was available in our lab. We gathered data on the mating system on 175 species: 105 that form pairs and 70 that do not. We obtained phenotypic information on the caregiving parent for 212 species: 86 species with biparental care and 126 that carry out maternal care.

To assess correlated evolution between traits, we used the ‘discrete method’ within the program BayesTraits (version V4.0; Pagel, 1994). Each trait was binarized for compatibility reasons. We run two different models: an independent model, where the two traits evolved separately, and a dependent model, where the evolution is correlated. We applied a Markov-Chain Monte Carlo (MCMC) with uniform priors to estimate Bayesian posterior distributions for rate parameters, and the ‘stepping stone’ sampler to obtain marginal log-likelihoods. To compare the models, we calculated the Log Bayes Factor (Log BF) as described in the manual. We used the reverse jump approach to assess precedence and contingency between the two traits (Pagel & Meade, 2006). We sampled a gamma prior with both parameters (shape and scale) ranging between 0 and 100, seeded from a uniform hyperprior ranging from 0 to 100. To test convergence, we applied Geweke’s convergence diagnostic in the Coda package (version 0.19-4.1) in R. For a detailed description of this section, see Supplementary Methods.

### 6. Tissue expression and correlation

To study the tissue expression of the nonapeptide precursors and receptors in Lake Tanganyika’s cichlids, we used the data from El Taher et al. (2021). Bulk RNA of six tissues (brain, liver, lower pharyngeal jaw, gills, ovaries and testis) of males and females of 73 species are available. We used the normalized count dataset available from El Taher et al. (2021). Using the brain counts, we then assessed correlation between gene expression levels of all nonapeptide system genes with pair-bonding behaviour and the sex of the caregiver. We performed a phylogenetic regression (Ives & Garland, 2010) for each gene. We implemented the phylogenetic regression with the function *phyloglm* (‘logistic_IG10’ method, 1000 independent bootstrap replicates, and p-values computed using Wald test) in the phylolm package (version 2.6.5). We included in the model data on stable isotopes linked to the diet (δ¹DN) and habitat (δ¹³C), obtained from Ronco et al. (2021). Because we could not include a sex variable in the formula of the function, we run an additional phylogenetic regression only with the individuals of each sex separately.

## Results

### 1. Nonapeptide system genes repertoire

Two nonapeptide precursors and eight receptors were found in the Nile tilapia genome (*Oreochromis niloticus*; RefSeq accession GCF_001858045.2, female). To assign a receptor type to each sequence, we built a phylogenetic tree with known gene sequences from distant teleost species. Based on sequence similarity with multiple teleost species, we assigned the following repertoire of genes in Nile tilapia: two *OTR*s, two *VTR1A*s, two *VTR2A*s and two *VTR2B*s (Supplementary Table 1). The species tree and sequences used can be consulted in Supplementary Data 1.

To find the gene repertoire in Lake Tanganyika cichlids, we performed a nucleotide BLAST search with the genes of the reference genome on all six long-read assemblies. Despite some genetic differences between Lake Tanganyika cichlids and the reference genome, we found a single clear match for each reference gene (e-value < 1e-200). Using the location on the Nile tilapia chromosomes, we compared the distribution of the genes across the long scaffolds in the six long-read assemblies (Supplementary Table 2). Both nonapeptide precursor genes were found in the same scaffold in a head-to-tail conformation and separated by 40-50 kbp, consistent with Nile tilapia’s annotation (Figure 1). The nonapeptide sequences in LT cichlids follow the canonical teleost OT (CYISNCPIG) and VT (CYIQNCPRG) sequences. The location of all nonapeptide receptors in the long-read assemblies was also consistent with the chromosomal distribution in Nile tilapia genome (Figure 1). For positive selection analysis, we selected splicing forms that differed to any extent in their protein-coding sequences. A total of 15 splicing forms across the 10 genes were taken from the reference genome, and one additional transcript was included from other cichlid annotations (Figure 1). Except for a 66 bp deletion in *VTR2Ab* gene, all genes showed a highly conserved protein-coding domain structure.

**Figure 1.**
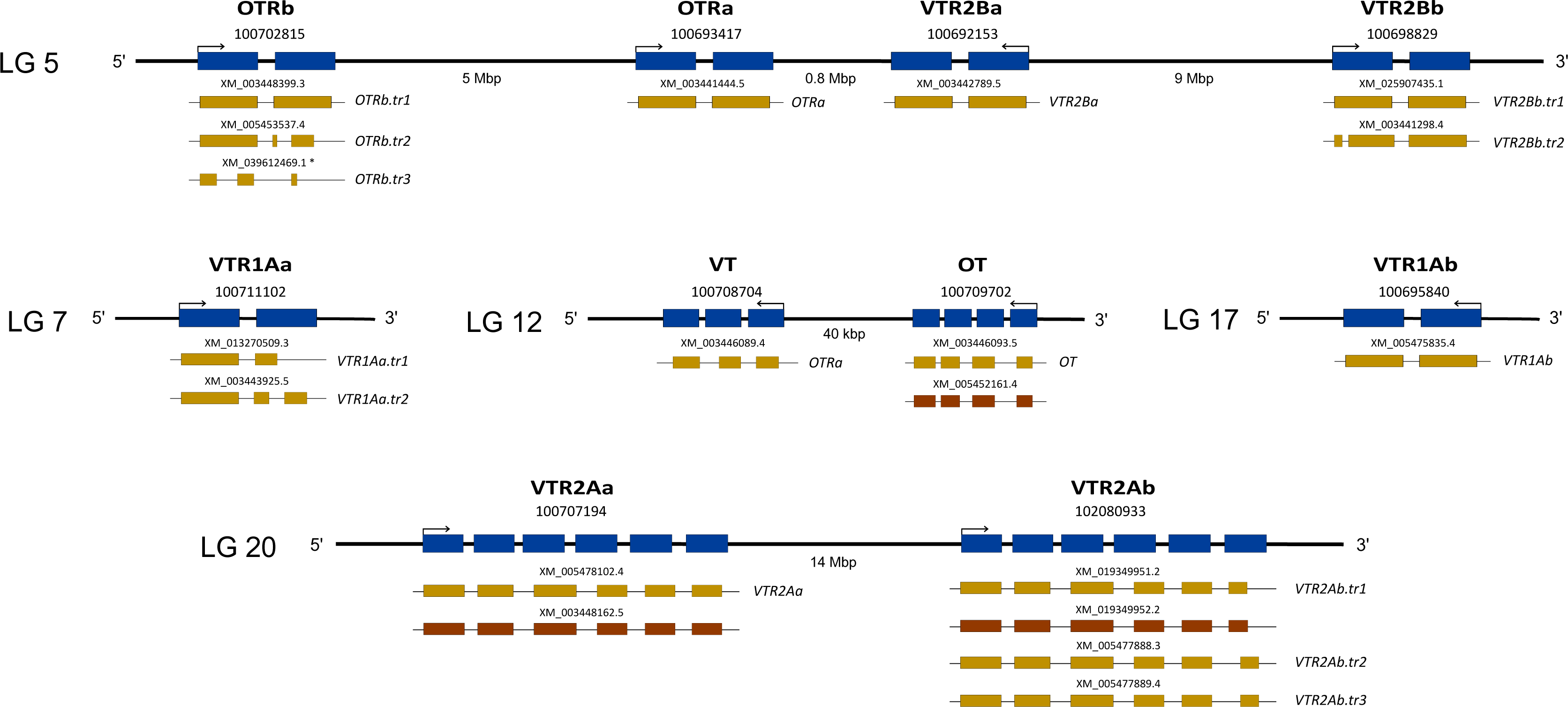
Genomic distribution of the nonapeptide system in the reference genome *Oreochromis niloticus*. In the present study we found evidence that suggests a similar distribution of the nonapeptide system genes in Lake Tanganyika cichlids. Gene IDs and transcript accession numbers are shown. All accession numbers were obtained from NCBI from Nile tilapia’s genome assembly GCF_001858045.2, except for *XM_039612469.1*. At the right of each transcript, the name of the transcript is given as it is referred to in this study. In blue, schematic representation of the exons comprising the 10 genes. In gold, transcripts selected for the present study. In brown, transcripts that did no differ from another transcript in their protein-coding sequence and therefore were not taken for analysis. The arrows indicate the sense of transcription. The genes are sorted by chromosome (linkage group, LG) number and location. (*) This transcript is not annotated in the reference genome, the accession number belongs to an *OTRb* transcript of another cichlid, the blue tilapia (*Oreochromis aureus*).

### 2. Structure of the nonapeptides and their receptors

The predicted nonapeptide precursors preproteins followed the known conserved structure that includes a signal peptide, hormone moiety, GKR residue, neurophysin and copeptin. All receptor genes had at least one isoform that exhibited the classic seven transmembrane (TM) domain structure, with three ILs and three ELs alternating between the transmembrane domains (Figure 2a). Some alternative splicing forms (*OTRb.tr1, OTRb.tr3 and VTR2Ab.tr3*) deviate from this canonical GPCR structure, resulting in shorter isoforms with fewer transmembrane domains and therefore fewer loops (Figure 2b, 2c, 2d). Additionally, we confirmed that the 66 bp deletion of *VTR2Ab* gene was located in the extended intracellular loop 3 (IL3). A detailed predicted domain structure of each transcript can be consulted in Supplementary Data 2.

**Figure 2.**
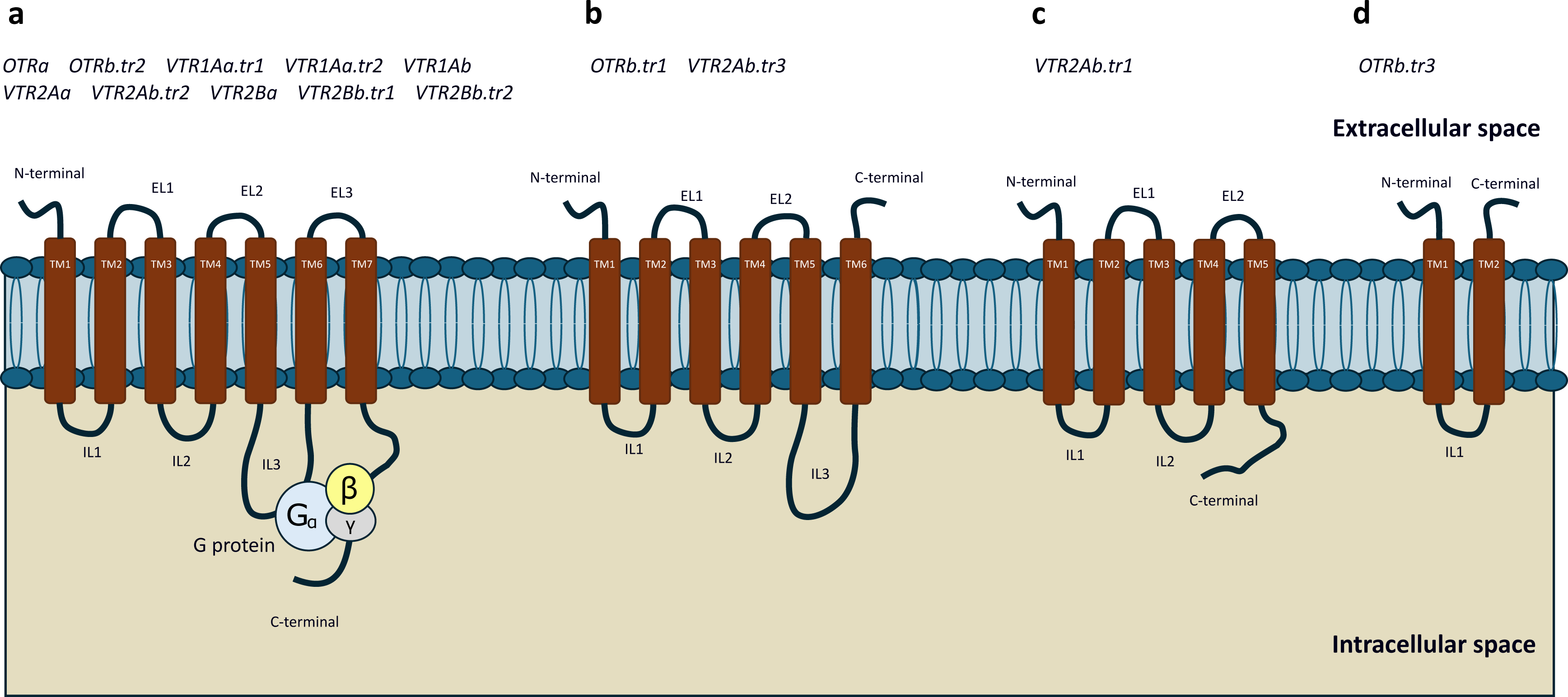
Schematic representation of the predicted domain structures of different alternative splicing variants of nonapeptide receptors in Lake Tanganyika cichlids. (a) Canonical G-protein-coupled receptor (GPCR) structure with seven transmembrane (TM) domains, three extracellular loops (EL1–EL3), and three intracellular loops (IL1–IL3), interacting with the G-protein complex. (b) *OTRb.tr1* and *VTR2Ab.tr3* isoforms, which lack the seventh TM domain, resulting in an extracellular C-terminus. (c) *VTR2Ab.tr1* isoform, consisting of five TM domains, two ELs, and two ILs. (d) *OTRb.tr3* isoform, which spans only two TM domains and has a single IL.

### 3. Nucleotide diversity of the nonapeptide system genes

*OTRa* exhibited the lowest nucleotide diversity (π) among all genes, followed by *OTRb*, *VTR1Aa*, and *VTR2Ba*. Both nonapeptide precursors displayed intermediate levels of variability, while *VTR2A* copies showed the highest diversity values (Supplementary Data 3).

Across domains, we only analysed the isoforms with a classic GPCR structure. Terminal regions appeared to be the most variable, while transmembrane domains and both extracellular and intracellular loops were less variable (Figure 3). Interestingly, the two extended IL3s (from *VTR2Aa* and *VTR2Ab* genes) showed the largest nucleotide diversity across all IL3. In addition, the C-terminal domains of both *VTR2A* copies are also among the most variable (Figure 3). The alignments and nucleotide diversity values can be consulted in Supplementary Data 3.

**Figure 3.**
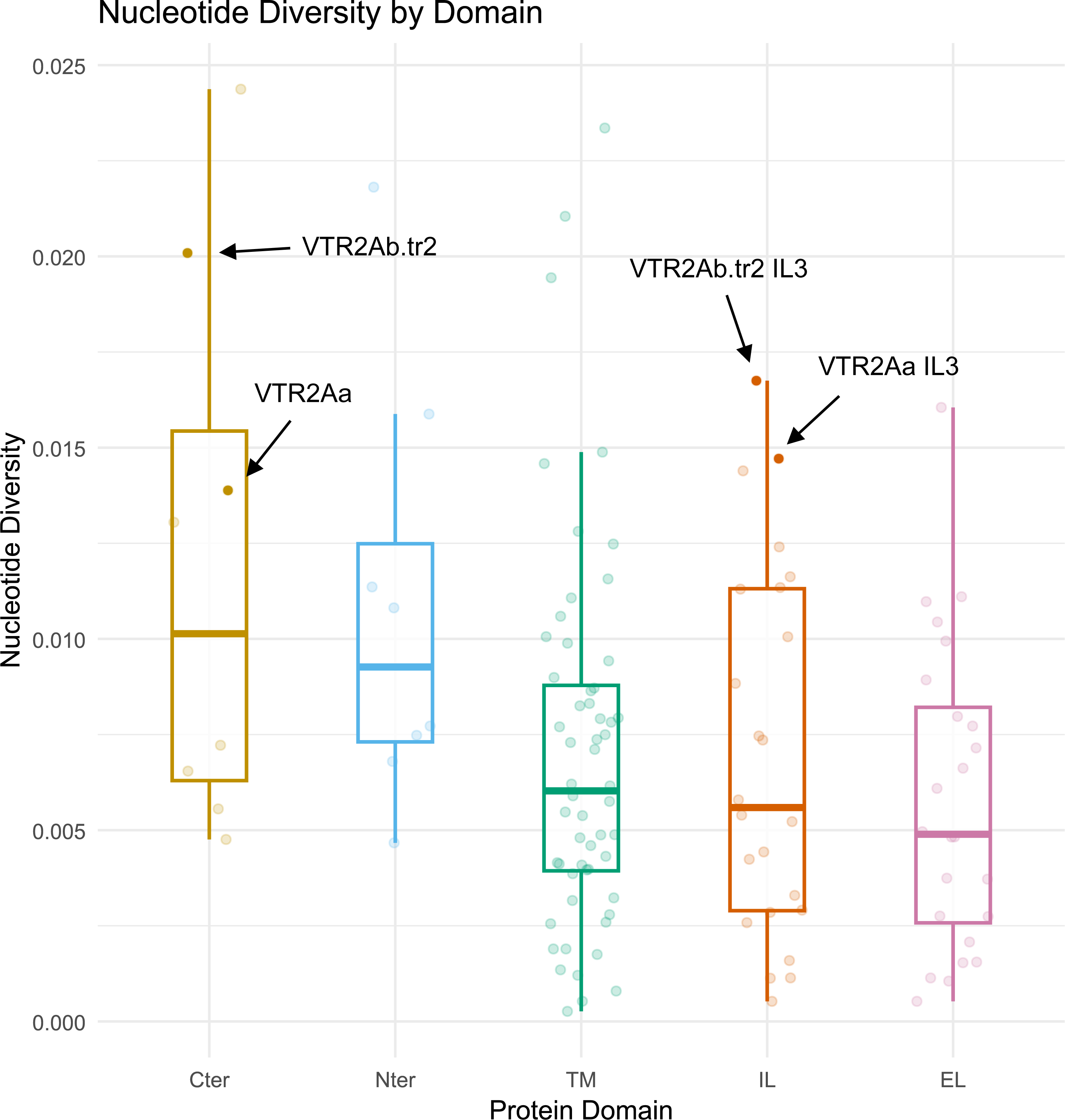
Nucleotide diversity (π) across protein domains of nonapeptide receptors. For each receptor gene, we selected one isoform with a predicted canonical GPCR structure. Abbreviations: Cter, C-terminus; Nter, N-terminus; TM, Transmembrane domain; IL, Intracellular loop; EL, Extracellular loop.

### 4. Positive selection on nonapeptide system genes

Gene-wide ω (dN/dS) values show generalized purifying selection across all genes. *OTRa* had the lowest ω, while both nonapeptides displayed relatively high values (Table 1). Both tree topologies yielded similar results. Similarly, both tree topologies provided largely similar results in site-specific positive selection analysis. The following description is based on the gene tree topology, as we considered that it better represents the evolutionary history of the gene. All genes and transcripts were subject to strong negative selection (Figure 4; Supplementary Data 4). A total of 335 sites under negative selection and 15 sites under positive selection (Supplementary Table 3). Across the receptors, almost half the negatively selected sites (42%) belonged to TM domains, followed by ILs (26%) and ELs (16%).

**Figure 4.**
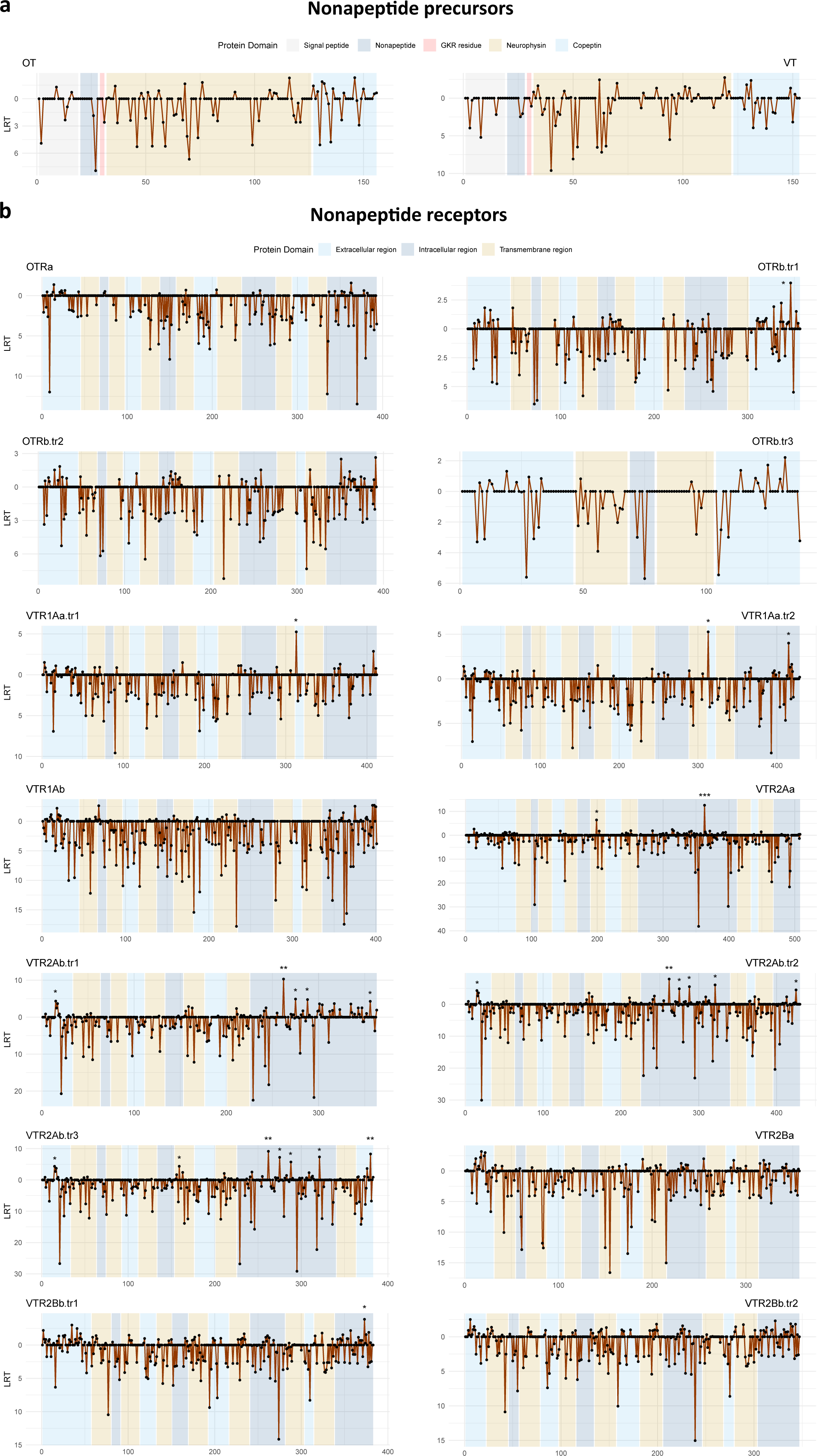
Likelihood ratio test (LRT) values of positive selection analysis along the protein sequences of nonapeptide precursors and receptors. X-axis represent amino acid number (site) along the protein sequence. LRT values represent the strength of selection at each codon position. For visualization purposes, LRT values are positive in both directions: upward deviations indicate positive selection, and downward deviations indicate negative selection. Each plot corresponds to a different transcript. Background shading represents protein domains. Asterisks indicate significance in positively selected sites(*) p-value < 0.05 (**) p-value < 0.005 (***) p-value < 0.0005.

No positively selected site was detected on the nonapeptide precursor genes. Similarly, *OTRa* showed no positively selected sites, and *OTRb* only one, unique to a specific isoform, located in the C-terminal extracellular region. Two positively selected sites were detected in *VTR1Aa* (one in EL3 and one in the intracellular C-terminal region), while *VTR1Ab* showed no positive selection. In contrast, *VTR2Aa* had two positively selected sites, with one site in IL3 being highly significant (p-value 0.0004). *VTR2Ab* exhibited the highest number of positively selected sites, with 6 in total, including four in the extended intracellular loop of its canonical copy *VTR2Ab.tr2* (Figure 5). In *VTR2Ba*, no positively selected sites were found under the gene tree topology. A detailed list of the sites under selection can be consulted Supplementary Data 4.

**Figure 5.**
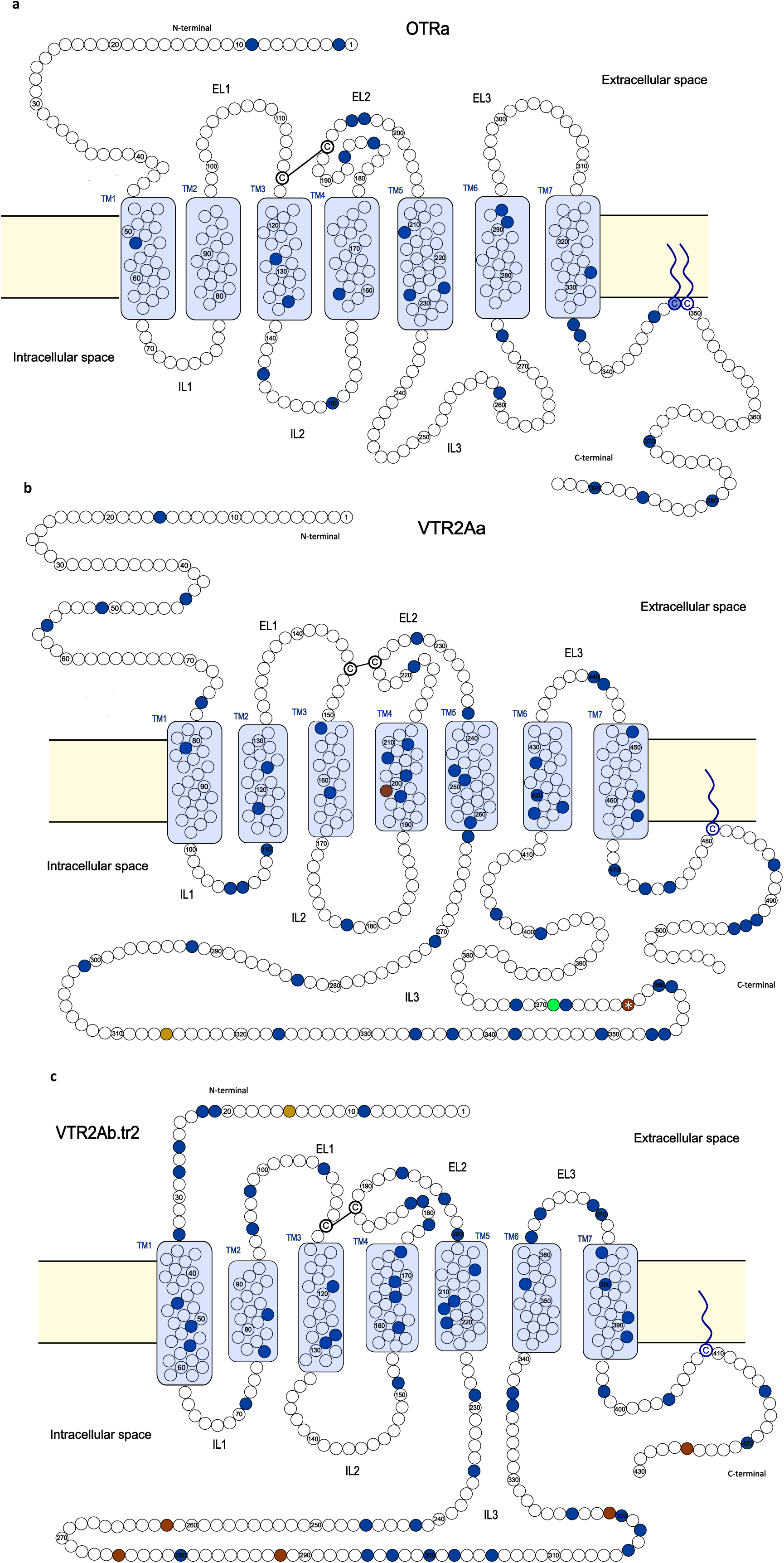
Schematic representation of nonapeptide receptor protein sequences showing selection analysis derived from dN/dS ratio: a) *OTRb*, b) *VTR2Aa*, c) *VTR2Ab.tr2*. Dark blue circles indicate significant negatively selected sites found with any of the two tree topologies (gene tree and species tree). Golden circles indicate significant positive selection found only with species tree (p-value < 0.05). Brown circles indicate significant positive selection found with both tree topologies (p-value < 0.05). Asterisk indicates very significant positive selection (p-value < 0.0005 when calculated with the gene tree topology). The green circle indicates an amino acid that showed correlated evolution with pair-bonding behaviour.

## Correlation between brooding and pair-bonding phenotypes

We found the two traits strongly correlated (log Bayes Factor 84.36) in Lake Tanganyika adaptive radiation (Figure 6). According to our reverse jump analysis, the ancestral state in Lake Tanganyika cichlids was a biparental species with a very high probability (∼92%), and very likely a pair-bonding species (∼74%). Together, the probability of a biparental and pair-bonding species was 70.42% The evolution of this binary traits seemed to follow a direction towards maternal and non-pair-bonding species. Pair-bonding behaviour seems to be easily lost when the caregiving is only maternal. Instead, gaining pair-bonding behaviour seems to be very unlikely (Supplementary Data 5).

**Figure 6.**
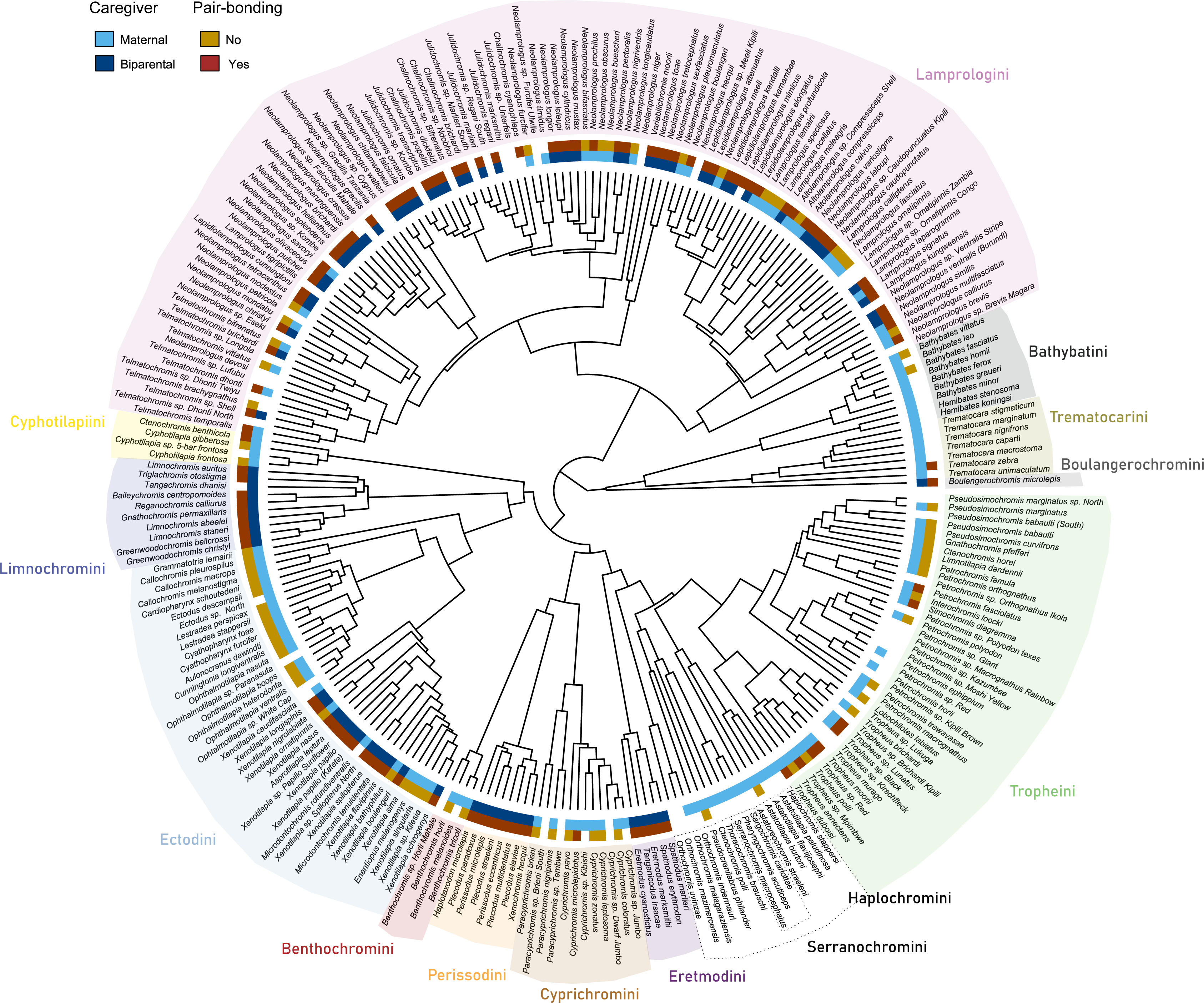
Phenotypic characterization of Lake Tanganyika cichlid radiation according to the two phenotypes used in the present study: sex of the caregiver and pair-bonding behaviour. Multiple phenotypic transitions can be observed. Background colors indicate tribe.

## Correlation between behavioural phenotypes and SNPs

Of all 1098 sites tested for correlation with both phenotypes, only one amino acid showed evolution in association with pair-bonding: position 369 of *VTR2Aa* transcript (green amino acid in Figure 5), with a log Bayes Factor of 2.5 (Supplementary data 5) with uninformative priors. In the reverse jump analysis, the three dependent models performed better than the independent with a log Bayes Factor of 6.67 (strong correlation, according to the manual), reinforcing that the correlated model had a better fit. This position is a valine-to-alanine substitution. The reverse jump analysis revealed a constraint with the amino acid alanine with pair-bonding species. Valine, the predominant amino acid at position 369, appears to accommodate both pair-bonding and non-pair-bonding states, whereas alanine is predominantly associated with pair-bonding species (Supplementary Data 5)

## Tissue expression and correlation with behavioural phenotypes

The nonapeptide precursors were almost exclusively expressed in the brain, their primary site of production. While *OTRa* was mainly expressed in the brain, suggesting a role in behaviour, *OTRb* showed very high expression in the gills. *VTR1Aa* was expressed in gills, lower pharyngeal jaw and testis, and to a lesser extent in the brain. The expression levels of *VTR1Ab* were generally lower and reduced to the gills and brain. All *VTR2*s were expressed in the brain (Figure 7). While the function of *VTR2s* in the brain remains less studied, their central expression appears to be consistent across teleosts (Lema et al., 2019; Martos-Sitcha et al., 2014). *VTR2Aa* expression was widespread across different tissues, suggesting a role in multiple physiological functions. Its high expression in the liver aligns with previous reports linking it to carbohydrate metabolism (Moon & Mommsen, 1990). Interestingly, its expression in the lower pharyngeal jaw was very high and its physiological role remains an open question. *VTR2Bb* showed expression in the ovaries and testis, in line with other studies (Rawat et al., 2019). The count data were extracted from El Taher et al. (2021). The count dataset used for this analysis can be consulted in Supplementary Data 6.

**Figure 7.**
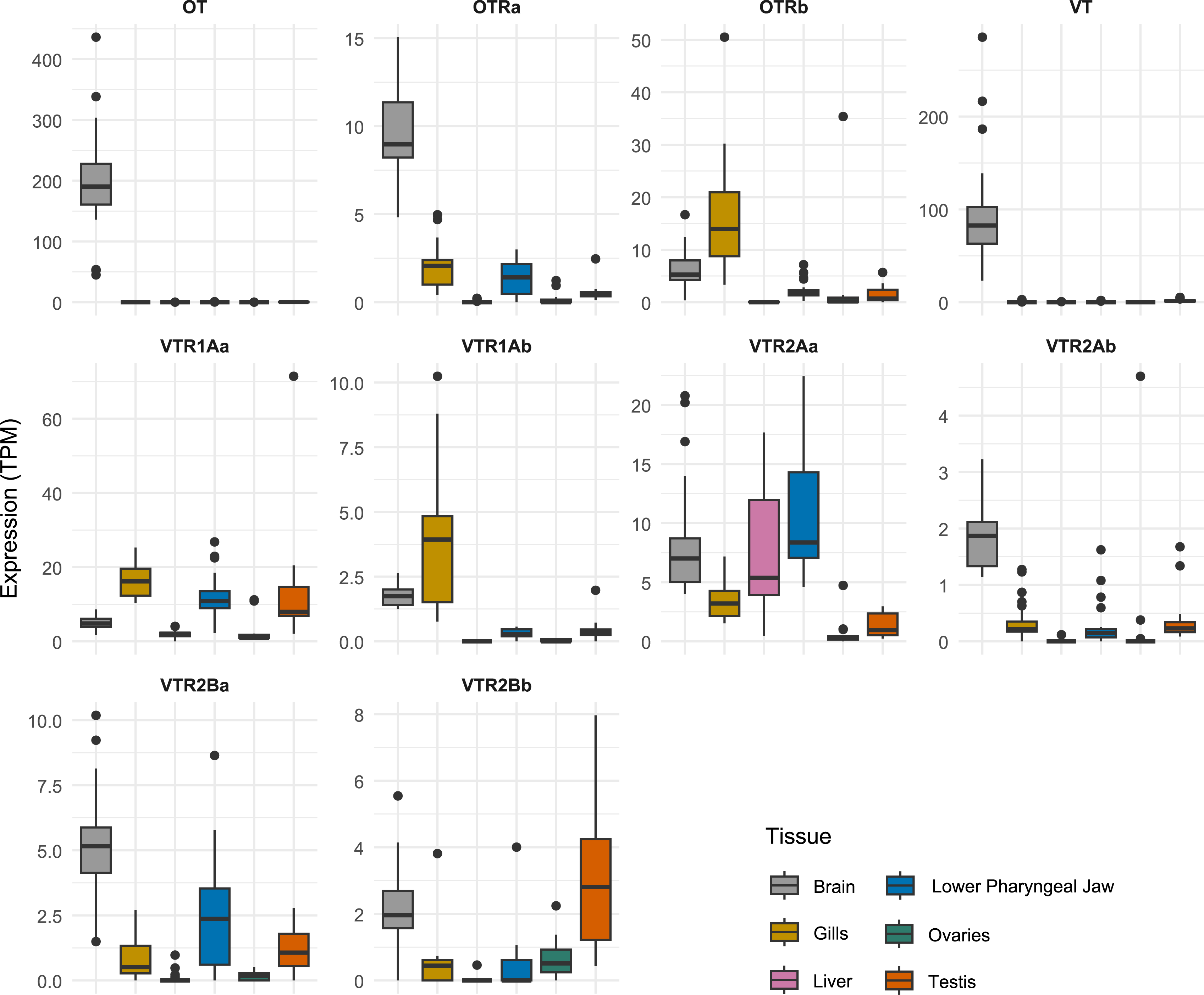
Tissue expression distribution of the nonapeptide system genes across the six genes. For visualization purposes, we excluded *Boulengerochromis microlepis* males from the nonapeptide precursor expression (*OT* and *VT*) because of exceptionally high expression levels. The values belong to whole tissue bulk RNA. TPM: Transcripts Per Million.

## Association between the expression of nonapeptide genes and behavioural phenotypes

Using a phylogenetic regression, we assessed if the expression levels of the nonapeptide system genes in the brain showed any statistical significance with pair-bonding behaviour and the sex of the caregiver. Although significance values alone should be taken with caution because of the nature of transcript count, comparing them provides insight into the potential role of gene expression levels in shaping phenotypic traits.

We observed multiple sex-specific differences. Regarding pair-bonding behaviour, we observed significant correlation with the phenotype in three genes: *OTRa*, *VTR1Ab*, and *VTR2Ab*. *OTRa* and *VTR1Ab* show higher expression in pair-bonding species, whereas *VTR2Ab* has higher expression in nonpair-bonding species. However, these correlations are only observed in males (Supplementary Data 4). When analysing the sex of the caregiver (maternal or biparental), we found that both *OTR* copies showed significant expression in males, with higher expression in biparental species. In females, significance was observed for *OT* and *VTR2Bb* expression, with biparental species showing higher expression (Supplementary Table 4).

## Discussion

We characterised the nonapeptide system gene repertoire in the adaptive radiation of cichlid fishes from Lake Tanganyika. We found that the Lake Tanganyika cichlids have one copy of each nonapeptide precursor in their genomes, and these are separated by 40-50 kbp. The nonapeptide sequences are the typical conserved teleost amino acid sequence of OT (CYISNCPIG) and VT (CYIQNCPRG). Our analysis based on long-read assemblies suggest that the receptor genes in Lake Tanganyika cichlids preserve the copy number and genomic arrangement found in the Nile tilapia. Despite minor genetic modifications, the overall exon structure is also conserved across Lake Tanganyika cichlids. Interestingly, *VTR2Ab* experienced a 66 bp deletion at an early stage in the radiation – after the divergence of Boulengerochromini, Bathybatini and Trematocarini lineages – that trims 22 amino acids of the IL3. A new early splicing site was acquired and the current widespread expression of this new variant evidences its functionality *OTRb* and *VTR2Ab* have alternative splicing forms whose predicted protein domain deviate from the canonical GPCR structure (7 TM domains alternating between 3 EL and 3 IL). The physiological function of the alternatively spliced receptor isoforms remains an open question. Notably, *OTRb.tr1* and *VTR2Ab.tr3* lack the final transmembrane (TM) domain, resulting in an extracellular C-terminus. This structural alteration has been reported to prevent receptor migration to the plasma membrane, retaining it within the ER-Golgi compartments (Gonzalez et al., 2011). Although all three extracellular loops contribute to ligand binding, EL1 and EL2 appear to be essential for nonapeptide-receptor interactions: EL1 binds the tripeptidic part of the nonapeptide, while EL2 interacts with the cyclic part (Gimpl & Fahrenholz, 2001; Postina et al., 1996; Waltenspühl et al., 2022). Therefore, OTRb.tr1, VTR2Ab.tr3 and VTR2Ab.tr1 may still retain their ligand-binding capacities, although potentially with altered affinity. In contrast, OTRb.tr3, which consists of only two transmembrane domains and a single intracellular loop, possesses extracellular domains limited to the N- and C-terminal regions. This structural configuration makes direct interaction with the ligand far less likely, raising questions about its functional role.

With the emergence of vertebrates and two associated whole genome duplication events, new copies of vasotocin receptors acquired new functions. The neofunctionalization of these receptors is very well established in mammals (Jurek & Neumann, 2018), but growing evidence are unravelling the implication of nonapeptides in a wide range of functions in fish (Balment et al., 2006), including social phenotypes such as aggression, pair-bonding or parental care in teleost (Godwin & Thompson, 2012; Nowicki et al., 2020; Reddon et al., 2017; Ripley & Foran, 2010). For neofunctionalization to occur, genetic diversification is necessary to introduce functional changes. To explore this possibility, we tested for positive selection in the nonapeptide system within the Lake Tanganyika cichlid adaptive radiation. We found that most amino acids were under strong negative selection, reflecting the strong evolutionary pressure to preserve protein structure and functionality. These negatively selected amino acids are distributed across all functional domains, consistent with the overall high conservation of the receptors across vertebrates (Gimpl & Fahrenholz, 2001; Mayasich & Clarke, 2016). Moreover, because of the young age of the radiation there are several sites that do not show any variation, which impedes making inferences about selection. 15 sites showed signs of positive selection when analysed with the gene tree topology. Interestingly, five of these sites were concentrated in the extended IL3 of *VTR2Aa* and *VTR2Ab*.

In our analysis, the two extended IL3 showed the largest nucleotide diversity among all intracellular loops. Furthermore, multiple or highly significant amino acids in this domain showed positive selection. Taken together, our findings suggest that the overall structure of *VTR2Aa* and *VTR2Ab* is under purifying selection, similarly to the other nonapeptide receptors, while the part in charge of signal transduction is subject to change. At some point in teleost evolution, the intracellular loop (IL3) of VTR2As expanded significantly while presumably retaining receptor functionality (Ocampo Daza et al., 2022). This enlargement introduced additional amino acids into the domain responsible for signal transduction, directly interacting with the G protein. While some residues remain highly conserved and under strong negative selection, reduced constraint in this extended region may allow for greater evolutionary flexibility, potentially facilitating functional diversification or adaptation to different regulatory mechanisms. This extended IL3 has drawn very little attention to date. However, IL3 is necessary and sufficient for activation of G_S_ (Liu & Wess, 1996), making it a crucial component of VTR2A downstream signalling. Further work should elucidate the functional implications and determine whether the extended IL3 contributes to specific regulatory mechanisms, signaling efficiency, or interactions with other intracellular partners. It is worth noting that our analyses of positive selection and interpretations primarily focused on the receptors that follow the canonical receptor domain structure. However, alternative variants may be subject to distinct selective pressures. Since they are transcribed from the same DNA sequence, functional divergence between isoforms could lead to selection acting more strongly on one, while changes in the other may occur as a byproduct.

In the cichlid adaptive radiation of Lake Tanganyika, multiple social phenotypes have repeatedly evolved, making it an ideal scenario to study the evolution of social phenotypes (Sefc, 2011). Our data suggest that the ancestral cichlid was a pair-bonding species that provided biparental care. In line with other studies (Goodwin et al., 1998; Iles & Holden, 1969; Klett & Meyer, 2002), we found that the evolutionary transition from monogamy and biparental care to non-pair-bonding and maternal care appears to be much more likely than the reverse. Therefore, biparental care seems to arise as a consequence of pair-bonding rather than serving as its initial driver. (Stanbrook et al., 2022) proposed that monogamy in cichlids arises for male mate guarding (Parker, 1970). This suggests that rather than evolving primarily to ensure biparental care, monogamy may have initially developed as a means for males to protect their mates from rival males, thereby securing their own reproductive success.

We hypothesised that the positive selection patterns we observed across the nonapeptide system could be linked to the evolution of social phenotypes. All receptors were expressed in the brain to some extent. The receptor is the place of action of the ligand, so we would expect that brain expression of the receptors is linked to behaviour. In order to assess if we would find correlates between nonapeptide receptors and social phenotypes, we first searched for correlation between the two social phenotypes (pair-bonding and sex of the caregiver) and the gene sequences.

While the sex of the caregiver did not show any correlation with any amino acid, a residue in the extended IL3 of *VTR2Aa* (position 369) showed evidence to be evolving in correlation with pair-bonding. The particular SNP results in a valine-to-alanine substitution. Valine has a larger side chain than alanine, which can influence the protein’s hydrophobic interactions, stability, and folding, potentially altering the protein’s activity. In addition, this amino acid is found only six amino acids downstream from the site with the strongest positive selection signal. The close proximity of this site to the most strongly selected residue raises the possibility that structural or functional modifications in IL3 of VTR2Aa play a role in modulating receptor signaling in the context of pair-bonding. Further functional studies would be necessary to determine the precise effects of this substitution on receptor activity.

Interestingly, although pair-bonding and the caregiver are strongly correlated, the VTR2Aa position 369 amino acid only showed correlation with pair-bonding but not the caregiver’s sex. This suggests that while pair-bonding and biparental care are linked, they may involve distinct genetic mechanisms. The amino acid variation in VTR2Aa may influence mate attachment without directly affecting parental care, indicating overlapping but separate molecular pathways.

We assessed correlation between gene expression and the phenotypes. The outcome of the phylogenetic regression should be interpreted carefully. While most genes did not show any statistical significance, we observed a lot of sex differences in nonapeptide system genes expression, which has been frequently reported (Albers, 2015; Dumais & Veenema, 2016). It is worth considering that expression in the brain does not necessarily correlate with its effect. In neural circuits, receptor abundance does not necessarily predict functional significance, as even low expression levels can influence signaling pathways and modulate behavioural responses. For this reason, especially with brain bulk RNA, positive signal can easily remain unnoticed. Finally, while previous research has documented VTR2A expression in various brain regions (Lema et al., 2019), its precise role in regulating behaviour remains largely unexplored. In our study, we observed relatively low expression levels of all VTR2s in the brain. However, low expression does not necessarily equate to low functional significance, particularly in neural circuits, where even subtle variations in receptor abundance can influence signaling pathways and behavioural outcomes. Given that our molecular evolution analyses suggest that the protein sequence of VTR2Aa is evolving in association with a social phenotype, it is reasonable to hypothesize that gene expression patterns may also be subject to similar evolutionary pressures. Further research integrating functional assays and expression analyses across diverse social contexts will be necessary to fully elucidate the behavioural roles of VTR2s in teleost.

## Conclusions

Our findings reveal that nonapeptide precursor and receptor genes are subject to strong purifying selection across the Lake Tanganyika cichlid radiation. While positive selection is rare, the few sites under selection are primarily concentrated in the extended intracellular loop of the vasotocin receptor 2A, suggesting potential functional diversification. Additionally, our correlation analyses highlight the involvement of nonapeptide receptors in shaping social phenotypes, reinforcing their role in the evolution of social behaviour in cichlids.

## Authors contributions

**Pol Sorigue**: Conceptualization; investigation; visualization; writing – original draft. **Walter Salzburger**: Conceptualization; writing – review and editing. **Rui Oliveira**: Conceptualization; Supervision; funding acquisition; writing – review and editing.

## Code availability

The scripts have been deposited on Github (https://github.com/psorigue/Molecular-evolution-OTVT-in-LT-cichlids).

## Acknowledgements

We thank Susana Varela for her valuable insights and help in the correlated evolution analysis using BayesTraits program. This study was funded by a grant from Fundação para a Ciência e a Tecnologia (FCT) awarded to RFO (PTDC/BIA-COM/3068/2020). PS is supported by a FCT PhD Fellowship. WS received funding from the Swiss National Science Foundation (208002).

