## Supplementary material for "Nonapeptide molecular evolution during the adaptive radiation of Tanganyika cichlids": Figure S1

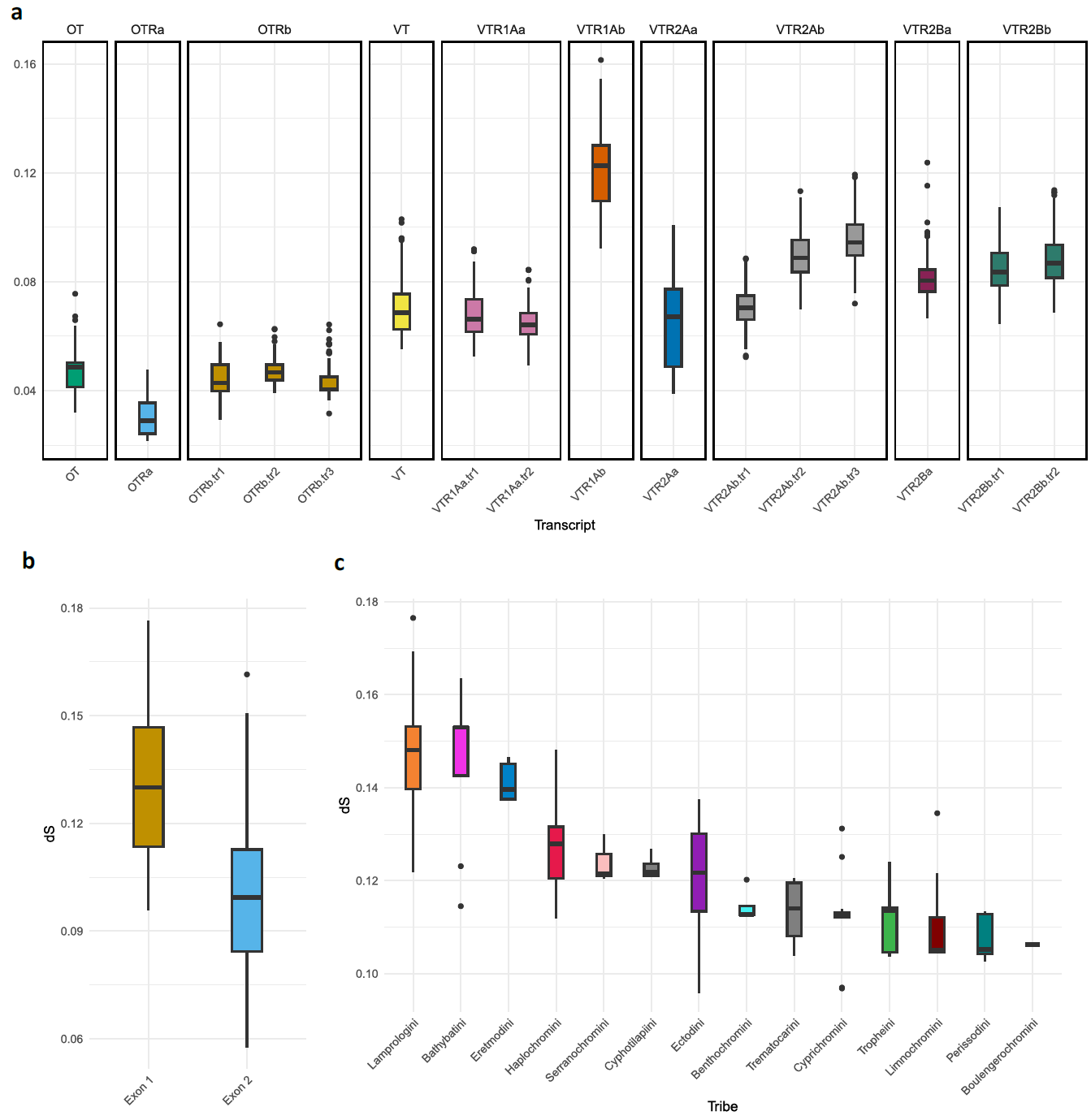

**Figure S1:** Rate of synonymous substitutions (dS) across protein-coding regions of the nonapeptide system. a) dS values for all transcripts. b) dS values of the two exons of VTR1Aa transcript. c) dS values of the first exon of VTR1Aa transcript across tribes.

The rate of synonymous mutations (dS) varied substantially across genes. Oxytocin-related genes exhibited lower dS values, with *OTRa* showing the smallest dS. In contrast, vasotocin and its receptors had higher values, with *VTR1Ab* standing out due to a substantially elevated dS compared to other genes (Figure S1). To investigate this further, we calculated dS values separately for the two exons of *VTR1Ab* and found that the first exon was responsible for the unusually high values (Figure S1). Additionally, when calculating dS by tribe, we observed that three tribes exhibited higher values: Bathybatini, Eretmodini and Lamprologini (Figure S1). Notably, Lamprologini, which accounts for almost half of the species in the radiation, had elevated dS values that significantly influenced the overall average. Given these high values, we considered the possibility of an overlooked factor. First, we recalculated dS using consensus sequences from PacBio assemblies, obtaining similar results. We then explored whether a hidden gene copy could be influencing the results. To test this, we performed a BLASTn search of the first exon to all six PacBio assemblies, but found no additional matches beyond the known gene. Although we cannot entirely rule out the presence of an undetected second copy masked by scaffolding limitations in the PacBio assemblies, we concluded that the higher dS values could be attributed to other biological factors or random variation rather than an undiscovered paralog. The dS values and the BLASTn results can be consulted in Supplementary Data 3.
