## Supplementary Methods for "Nonapeptide molecular evolution during the adaptive radiation of Tanganyika cichlids"

**Lake Tanganyika cichlids *de novo* assemblies**

The Illumina whole-genome raw reads from all cichlid species in Lake Tanganyika (Ronco et al., 2021a) were downloaded from the Sequence Read Archive (<https://www.ncbi.nlm.nih.gov/sra>). *De novo* assemblies were created for all available individuals. Genomic raw reads (Bioproject PRJNA550295; NCBI BioProject database) were mapped to the reference genome using Bwa-mem (version 0.7.18; (Li & Durbin, 2009) with default parameters. In Bioproject PRJNA550295, genomes for all the species of the radiation are available, ranging from 1 to 4 individuals per species. Transcriptomic raw reads (Bioproject PRJNA552202(El Taher et al., 2020) were mapped to the reference genome using STAR aligner (version 2.7.10a; (Dobin et al., 2013) with the options *outFilterMultimapNmax 1*, *outFilterMatchNminOverLread 0.4*, and *outFilterScoreMinOverLread 0.4*. All six tissues available (brain, gills, lower pharyngeal jaw, liver, ovaries and testis) and 4 to 6 individuals across 73 species were used for the analysis. We kept genomic *de novo* assemblies for each individual. However, to reduce complexity and increase the species coverage, we merged the transcriptomes of all the individuals of the same species into one species assembly for each tissue.

**Nonapeptide system in Nile tilapia**

For the present study we used Nile tilapia (*Oreochromis niloticus*) as the reference genome. Nile tilapia is an African cichlid phylogenetically equidistant to Lake Tanganyika’s adaptive radiation that has proven to be a useful reference genome in previous studies (El Taher et al., 2021; Ronco et al., 2021b). To identify the full repertoire of nonapeptide precursors and receptor genes in Nile tilapia, we used two widely studied teleost species: zebrafish (*Danio rerio*) and medaka (*Oryzias latipes*). We obtained the protein sequences of the nonapeptides (Banerjee et al., 2017) and their receptors (Ocampo Daza et al., 2022), and we searched similar protein sequences using protein BLAST (“BLASTp”, <https://blast.ncbi.nlm.nih.gov>) in our reference genome (*Oreochromis niloticus*; RefSeq accession GCF_001858045.2, female). All tilapia’s sequences that matched the search (e-value < 1e-50 ) were kept for further analysis.

To identify the receptor types, we built a phylogenetic tree with the protein sequences of representatives of distant branches within the teleost clade: Zebrafish (*D. rerio*), Medaka (*Oryzias latipes*), Eastern happy (*Astatotilapia* *calliptera*), and Electric eel (*Electrophorus electricus*). We first aligned the sequences with MAFFT (version v7.526; Katoh & Standley, 2013) and then we built a maximum likelihood phylogenetic tree using IQTREE (version 2.0.3; (Nguyen et al., 2015):. We included the human (*Homo sapiens*) receptor proteins and the Common octopus (*Octopus vulgaris*) octopressin receptor sequence as an outgroup. The accession numbers of all the sequences can be consulted in Supplementary Table 1.

We assigned the type of receptor by sequence similarity, that is proximity in the phylogenetic tree. The final repertoire of Nile tilapia’s nonapeptide system consisted of ten genes: two nonapeptide precursors – OT and VT – , two OTRs – OTRa and OTRb –, and six VTRs – VTR1Aa, VTR1Ab, VTR2Aa, VTR2Ab, VTR2Ba and VTR2Bb –. No VTR2C copy was found.

**Nonapeptide system in Lake Tanganyika cichlids**

We investigated the presence and number of copies of all nonapeptide precursor and receptor genes in Lake Tanganyika cichlids. To this end, we used long-read PacBio assemblies (of six cichlid species representing six tribes: *Neolamprologus multifasciatus* (Lamprologini; GenBank assembly GCA_963576455.2, *Simochromis diagramma* (Tropheini; GenBank assembly
GCA_900408965.1), *Bathybates minor* (Bathybatini), *Cyphotilapia frontosa* (Cyphotilapiini), *Cunningtonia longiventralis* (Ectodini), and *Cyprichromis leptosoma* (Cyprichromini) (unpublished). We created a local database with each long-read assembly using ‘makeblastdb’ (version 2.16.0+; <https://www.ncbi.nlm.nih.gov/books/NBK569841/>). This allowed us to search sequences in the database matching our query sequences. We then searched for the presence of all genes in each long-read assembly. To this end, we performed nucleotide BLAST (“BLASTn”) searches with the reference gene sequences as query. In all six long-read assemblies we found a clear match for each gene spanning the entire reference gene (e-value < 1e-200), although a second copy of VTR1Aa was found in *N. multifasciatus* assembly (discussed below). Small fragments of some genes had multiple matches along the genome. We concluded that this repeated matching was due to shared domains with other proteins unrelated to the nonapeptide system. The location of the nonapeptide system repertoire in the six long-read assemblies can be consulted in Supplementary Table 2.

Because dN/dS-based positive selection analyses are based on protein-coding sequences, we searched for protein-coding sequences in the reference genes in the Nile tilapia’s RefSeq annotation. We analysed all the annotated splicing forms of all 10 genes, and downloaded the coding sequences of all the isoforms that differ to any extent in their protein sequence.

To ensure a complete set of splicing forms produced in Lake Tanganyika cichlids, we did a protein BLAST search with the term “African cichlids” with all the nonapeptide system genes. In combination, we also manually inspected the mapped brain transcriptomes in Integrative Genome Viewer (IGV, version 2.16.1) to spot expressed isoforms that are not annotated in Nile tilapia’s assembly. Only one additional isoform of *OTRb* (later named *OTRb.tr3*) was robustly expressed among Lake Tanganyika cichlids and therefore added in the analysis. This transcript appears frequently expressed among the brain transcriptomes, and is annotated for other African cichlids such as *Oreochromis aureus*, *Astatotilapia burtoni* and *Neolamprologus brichardi* (accession numbers XM_039612469.1, XM_042223042.1, and XM_035904568.1, respectively).

Selecting only those genes that differed on their coding sequences, only one copy of each ligand precursor gene was found, while multiple transcripts were found for some receptors. A total of 16 splicing forms across the 10 genes were kept for further analysis. For a complete list of accession numbers, coding sequences locations and transcript names, see Supplementary Table 1. In the case of genes with multiple isoforms, we named the transcripts using the gene name followed by ‘tr’ and a number. This number was assigned based on the order in which the transcripts were identified during the analysis, without following any specific criteria. The genes with only one single transcript, this was named with the gene name.

To extract the consensus sequence of each species and gene, we used the newly assembled Illumina genomic and transcriptomic reads. We first assessed whether both genomic and transcriptomic assemblies contained reads mapped at the gene locations. A bad coverage can lead to not representative consensus sequences that eventually can strongly bias the analysis of positive selection. The coverage was analysed using Samtools depth (Danecek et al., 2021). This provided a per-site value corresponding to the number of reads mapped on the particular location. Combining manual inspection of the BLASTn results obtained from the long-read assemblies and the Illumina assemblies coverage files (Supplementary Data 3), we identified the following genetic features that warranted further investigation.

1. **Second copy of *VTR1Aa* in *N. multifasciatus***

The BLASTn search of Nile tilapia’s *VTR1Aa* to the long-read *N. multifasciatus* assembly resulted in two very similar copies found in different contigs. One of the copies presented multiple insertions and one deletion in the second exon, presumably disrupting the functional reading frame.

To investigate whether other Lamprologini species also possess a second *VTR1Aa* copy, we designed an artificial reference sequence based on *N. multifasciatus* 10 genes extracted from its long-read assembly including the duplicated copy of VTR1Aa in our reference sequence. To prevent intergenic splicing and ensure that mapped reads would only align to the intended regions, we flanked each gene sequence with 10000 ‘N’ nucleotides on both sides. We then mapped the raw transcriptomic reads from all available Lamprologini species onto this artificially constructed sequence to determine whether both copies of VTR1Aa were transcribed across multiple species. We used STAR aligner with very stringent parameters (*outFilterMultimapNmax 1*, *outFilterMatchNminOverLread 0.9*, and *outFilterScoreMinOverLread 0.9*), to avoid multimapping or erroneous mapping .To quantify the expression levels of each *VTR1Aa* copy, we employed Samtools coverage (version 1.21) to measure the depth of read coverage across the two gene copies. If both copies were present and actively transcribed in multiple Lamprologini species, we would expect to observe comparable or significant coverage at both loci. In contrast, if the second copy were unique to *N. multifasciatus*, the transcriptomic reads from other species would likely only map to the original VTR1Aa locus, showing little to no coverage at the duplicated region.

2. **Frameshift in OTRa first exon in *S. diagramma***

*S. diagramma* long-read assembly showed a frameshift due to an insertion of one nucleotide in the first exon of *OTRa* protein-coding region. This change in the reading frame completely changes the predicted protein product and therefore it would become not functional. To investigate this further, we visualized the mapped reads of this species’ Illumina assemblies (both genomes and transcriptomes). In both cases, we saw a correct alignment of the reads and no frameshift was observed. We concluded this could be due to a technical error during the sequencing or assembling, or due to an individual singularity

3. **Sorter fourth exon of VTR2Ab**

Except for the three most ancient clades (*Boulangerochromini, Trematocarini, Bathybatini*), we observed in all species a region within *VTR2Ab* gene with no coverage. We manually inspected the mapping files with a genome viewer (IGV) and spotted a 66 bp deletion at the C-terminus of the fourth exon of *VTR2Ab*, just before a splicing junction. This exon is transcribed in all three splicing forms of the gene.

To evaluate the impact of this deletion, we examined transcriptome BAM files to identify reads supporting functional splicing. Specifically, we analysed the junctions file generated by STAR mapping, which records the precise locations where introns have been excised during pre-mRNA splicing. We searched for junctions that would fall at the neighbouring regions of the deletion. Multiple species across different tribes showed evidence of an isoform that is expressed and keeps the correct reading frame, although reducing the sequence length with an early splicing site that follows the canonical motif GT/AG. Given the widespread conservation of this new splicing across the radiation, we decided to trim the deletion and keep the shorter isoform for further analysis. The deletion location is NC_031984.2:6394290-6394355 in the reference genome and likely occurred after the divergence of the clades Bathybatini, Boulangerochromini, Trematocarini. The coverage file can be consulted for the full and trimmed versions of the three transcripts of *VTR2Ab* in Supplementary Data 3.

4. **Deletion of 16 bp in the sixth exon of VTR2Ab**

*VTR2Ab.tr3*, has a sixth exon that comprises the last 12 codons of the protein and the 3’UTR. However, several Lamprologini species have a 16 bpdeletion in the middle of this exon. In multiple species, we observed both haplotypes in the same species. The deletion presumably changes the reading frame. However, given that is standing variation, we decided to include this fragment in the analysis. The genomic location of this deletion in the reference genome is NC_031984.2: 6395547-6395562.

5. **Partial copy of VTR2Bb**

Across multiple long-read assemblies (*Simochromis diagramma*, *Cyphotilapia frontosa*, *Cunningtonia longiventralis*) we observed a partial tandem duplication upstream of the first exon of VTR2Bb. However, we observed changes in the reading frame caused by insertions and deletions along the duplicated copy. Therefore, we concluded that this copy is not functional and it is just the result of a segmental duplication that may have no impact on VTR2Bb protein product.

**Consensus sequences**

The consensus sequence of each gene was obtained for each individual using the genomic location of the coding sequences and the mapping files (BAM). A consensus sequence was obtained from the genomic mapped reads for each individual. By merging the individual sequences with Seqtk (<https://github.com/lh3/seqtk/>), we further obtained a species consensus. In the species consensus we kept intraspecific variation with ambiguous nucleotide symbols. The sequences were merged using In addition, we extracted a consensus from the transcriptomic mapped files. In this case, only a species consensus was obtained from all available individuals of each species. To obtain the consensus from the BAM file, a combination of SAMtools and BCFtools was used (Danecek et al., 2021). All consensus sequences can be consulted in the multiple sequence alignment (MSA) files in Supplementary Data 3.

**Multiple Sequence Alignments**

We built a multiple sequence alignment for each gene with the consensus sequences of all Lake Tanganyika cichlids and the reference gene from Nile tilapia genome. To improve the accuracy of the species representative sequences, we used both genomes and transcriptomes consensus. For each gene, we computed the number of nucleotides that had no mapped read for each species and each source (genomes and transcriptomes). Then, for each species we used the consensus sequence with less unmapped positions. In the vast majority of the genes and species, the genomic consensus featured better coverage (Supplementary Data 3). We performed a multiple sequence alignment using MAFFT (version v7.526; (Katoh & Standley, 2013). We then translated every coding sequence into amino acids, and checked for the presence of START/STOP codons, early STOP codons or sequence lengths not dividable by 3. We spotted multiple STOP codons with the sequence ‘TGA’. Although the codon ‘TGA’ is known as a translation STOP sequence, this also encodes a selenocysteine and does not necessarily stops the translation (Rajput et al., 2019). However, this codon is commonly taken as a STOP codon in computational programs (as it is the case in PAL2NAL and Hyphy used in the present study), thus not allowing downstream analyses. To make it compatible with the programs, we substituted nonterminal TGA codons by ‘NNN’. This was only the case in VTR2Ab.tr1 and VTR2Bb.tr1. In addition to this, the following manual changes were done on the alignments:

- ***VT***: Lake Tanganyika cichlid species do not have a start codon (‘ATG’) in the same location as the reference *VT* gene. In addition, some species have a STOP codon after the reference START codon. For this reason, we presumed the start of VT protein 11 amino acids downstream, where another ‘ATG’ is found. Therefore, we trimmed these 11 amino acids from the alignment.
- ***OTRb.tr1***: Six Lamprologini species have an early STOP codon: *Lamprologus kungweensis, Lamprologus laparogramma, Lamprologus* sp. "Ornatipinnis Congo", *Lamprologus ornatipinnis, Lamprologus* sp. "Ornatipinnis Zambia", and *Lamprologus signatus.* This STOP codon is found 40 amino acids to the end, and only affects the C-terminal intracellular domain. Given that we cannot directly assess the functional impact of this fragment, we decided to trim the downstream fragment that falls after the early STOP site only in the species where this was the case.
- ***OTRb.tr3***: In many species we found an early STOP codon that was found two amino acids upstream. We concluded that the protein can still be functional and we trimmed these amino acids in all the cases where the early STOP codon was found.
- ***VTR2Bb.tr1***: Several species showed a frameshift located after the reference START codon, which disrupted the entire reading frame of the gene in Lake Tanganyika species. Because we found another ‘ATG’ 12 amino acids downstream of the reference START site, we trimmed the initial 12 amino acids and presumed a new START site.

After the manual modifications, a codon-based alignment was then performed using PAL2NAL (version V14; (Suyama et al., 2006) in the command-line. With this step we ensured that positive selection analysis was done on the correct reading frame. The resulting alignments were the starting point for all the following analyses. The

**Structure**

To assign each amino acid to a domain of the protein, we used InterPro (<https://www.ebi.ac.uk/interpro/search/sequence/>), a comprehensive database for predicting protein domains based on amino acid sequence. Using this tool, we obtained information on amino acid positions and domain architecture location along the peptide sequence.

**Nucleotide diversity**

We calculated the nucleotide diversity index (π) of the alignments using the Ape package (version 5.8) in R (version 4.4.0). This index quantifies the average number of nucleotide differences per site between all the pair-wise combinations of species sequences, and provides one value for the entire alignment. Higher values indicate more genetic diversity between the sequences.

To examine domain-specific variation, we calculated π for all domains in each canonical isoform. We selected both ligand transcripts and only the receptor transcripts that are predicted to have seven TM domains that alternate between 3 intracellular loops and 3 extracellular loops, and flanked by an extracellular N-terminus and intracellular C-terminus. To avoid biasing the nucleotide diversity values, we selected only one isoform for each receptor gene. We first calculated nucleotide diversity (π) for each of the individual domains. In nonapeptide precursor this included 4 domains per gene: signal peptide, nonapeptide, neurophysin and copeptin. In the case of the receptors, a total of 15 individual domains were calculated: seven transmembrane (TM1 to TM7) domains, three extracellular loops (EL1 to EL3), three intracellular loops (IL1 to IL3), N-terminus, and C-terminus. Finally, we computed π by grouping domains by type to determine whether certain regions of the protein exhibit more variation than others. This approach helped us assess whether specific structural features influence sequence diversity across different domain types.

**dS**

We analysed the rate of synonymous substitutions (dS) across species for all 16 transcripts. Comparing dS across different genes and lineages helps assess whether these genes are evolving under evolutionary constraints. Additionally, in genomes that are still assembled at the contig level, variation in dS can provide evidence for the presence of hidden gene copies. While the accumulation of synonymous substitutions is often used as a molecular clock, we rely on a reference genome that is evolutionarily equidistant from all Lake Tanganyika cichlid species. Therefore, we do not expect an excess of time-driven synonymous mutations. We calculated the rate of synonymous substitutions for all transcripts and all species using Yang and Nielsen method (Yang & Nielsen, 2000).

We obtained a particularly high value of dS in VTR1Ab gene. We then calculated the dS of the two exons separately, and we observed that the source of high dS was the first exon. We explored if there was a second copy of this exon in the genomes. For this reason, we used BLASTn to find hits of the first exon of VTR1Ab in the six long-read assemblies. However, in none of the six assemblies we found any duplicate of the first exon of VTR1Ab.

**Gene trees**

For each transcript, we built a ML phylogenetic tree using IQTREE (version 2.0.3; (Nguyen et al., 2015). Among different substitution models, IQTREE automatically selects the best fit and creates a phylogenetic tree. We used nucleotide and amino acid sequences of each transcript to build the trees.

**Positive selection**

Whole-gene ω (dN/dS) values were obtained using HyPhy (version 2.5.62) with the Fixed Effects Likelihood (FEL) method ((Kosakovsky Pond & Frost, 2005) using both tree topologies (gene tree and species tree). We also used FEL to investigate the occurrence of positive selection across the gene sequences of the nonapeptide system genes. It is commonly assumed that the rate of synonymous substitutions (dS) is neutral, and many programs designed to detect positive selection operate under the assumption that dS remains constant across all lineages. However, in our dataset we saw large variability between species. Moreover, allowing varying dS has been shown to improve accuracy (Pond & Muse, 2005). We selected FEL as an appropriate method because it permits variation in dS across both sites and branches. We accepted a p-value of < 0.05 as a threshold of significance.

**Phenotypic characterization**

We gathered phenotypic data for all species for which information was available in our lab. We gathered data on the mating system on 175 species: 105 that form pairs and 70 that do not. We obtained phenotypic information on brooder’s sex for 212 species: 86 species with biparental care and 126 that carry out maternal care. We classified species as pair-bonding if mates remained together beyond the act of reproduction, maintaining a social or physical association after mating. Facultative monogamous species were considered as ‘yes’ in pair-bonding. Species were considered biparental if males contributed directly to offspring care. However, territorial defense alone, without direct involvement in nurturing or protecting the young, was not considered a form of brood care. The phenotypic data for all species can be consulted in Supplementary Data 5.

**Correlated evolution**

We used the ‘discrete method’ within the BayesTraits (version V4.0; (Pagel & Meade, 2006) program to test for correlated evolution between pair-bonding (either ‘yes’ or ‘no’) and the sex of the caregiver (either ‘biparental’ or ‘maternal’) along the species tree (Ronco et al., 2021a). Each trait was binarized for compatibility with the program. We run two different models in BayesTraits discrete method: an independent model, where the two traits evolved separately, and a dependent model, where the evolution is correlated. Following the manual information, we chose a uniform prior, as it makes the fewest assumptions. We applied a Markov-Chain Monte Carlo (MCMC) with uniform priors to estimate Bayesian posterior distributions for rate parameters, and the ‘stepping stone’ sampler to obtain marginal log-likelihoods to compare models. To compare the models, we calculated the Log Bayes Factor (Log BF) as described in the manual:

Log BF = 2*(log marginal likelihood complex model – log marginal likelihood simple model)

A Log BF greater than 2 provides support for the correlated model, in which the variables evolve together ((Pagel & Meade, 2006).

We used the reverse jump approach to assess precedence and contingency between the two traits. This method forces evolutionary transition rates to be 0 in other to later compare free models against restricted models. We sampled a gamma prior with both parameters (shape and scale) ranging between 0 and 100, seeded from a uniform hyperprior ranging from 0 to 100. All commands used in BayesTraits are shown in Supplementary Data 5. We assessed convergence of the MCMC sampling using the Coda package (version 0.19-4.1) in R. Specifically, we applied Geweke’s convergence diagnostic, which compares the first 10% and the last 50% of the iteration chain to determine whether convergence has been achieved. We considered values between -1.95 and 1.95 indicative of good convergence, and any sites that did not meet this threshold were reanalysed with additional iterations.

We applied the same method to assess correlated evolution between a trait and amino acid information. For each amino acid position in our alignments that showed variation, we binarized the two most frequent alleles at that position. In case of ambiguous amino acids we included them as missing data. In cases where a species possessed a third or more alleles, the corresponding data points were removed from the dataset. Then we assessed correlation between the allele and one of the traits at a time. We discarded the sites in which the minor allele was present in less than 5 species. A total of 1098 sites were analysed independently following the same workflow explained for association between traits. In case of correlated evolution, we run again the analysis 100 times to ensure a stable likelihood. In addition, we randomized the SNP variable using the *sample* function in R, keeping the counts of each major/minor allele, and run correlation analysis with the randomized datasets. With this, we assessed whether the observed correlations were robust to randomization, ensuring that any detected association was not due to chance. By repeating the analysis with randomized SNP data, we could determine whether the likelihood values and inferred correlations were significantly different from those expected under a null distribution, thereby strengthening the validity of our findings.

**Tissue expression**

To study the tissue expression of the nonapeptide precursors and receptors in Lake Tanganyika’s cichlids, we used the data from (El Taher et al., 2021). In this study, they extracted bulk RNA of six tissues (brain, liver, lower pharyngeal jaw, gills, ovaries and testis) of males and females of 73 species. We used the normalized count dataset available in this study’s publication. We extracted the counts of the nonapeptide precursor and receptor genes from the dataset by filtering based on gene IDs. We computed the median expression level across replicates for each gene. All expression analyses presented in this study were conducted using the dataset available in Supplementary Data 6.

**Phylogenetic regression**

We assessed correlation between gene expression levels of all nonapeptide system genes with pair-bonding behaviour and the sex of the caregiver. Expression level usually has a strong phylogenetic signal (El Taher et al., 2021) that will bias the results if it is not considered. For this reason, to find correlates between gene expression and the phenotypes we used a phylogenetic logistic regression as described in (Ives & Garland, 2010). The phylogenetic regression runs a linear model that accounts for phylogenetic distance. We implemented the phylogenetic regression with the function *phyloglm* (‘logistic_IG10’ method, 1000 independent bootstrap replicates, and p-values computed using Wald test) in the phylolm package (version 2.6.5). Gene expression usually has large variability that is dependent on many factors. In our attempt to reduce this variability, we included in the model data on stable isotopes linked to the diet (δ¹⁵N) and habitat (δ¹³C), obtained from (Ronco et al., 2021). The function *phyloglm* requires one data point per species, so we computed the median of normalized gene counts of all the replicates for each species. We standardized gene counts, and isotope data. Because only one data point for species can be taken by the function, we could not include a sex variable in the function. For this reason, we run an additional phylogenetic regression only with the individuals of each sex separately.
