## Supplementary figures and images for "Nonapeptide molecular evolution during the adaptive radiation of Tanganyika cichlids"

### dS_all_genes.pdf

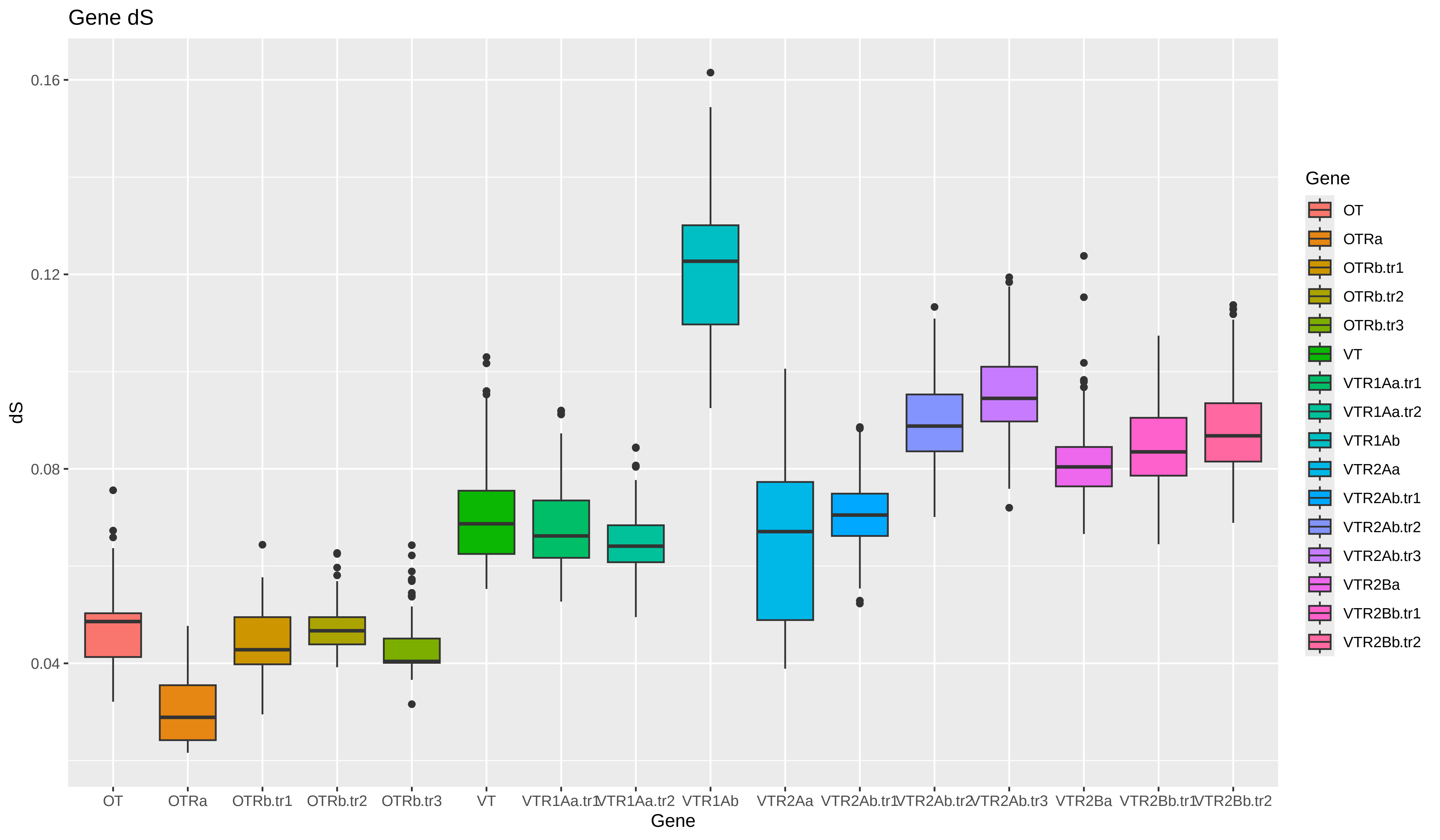

### dS_by_exon.pdf

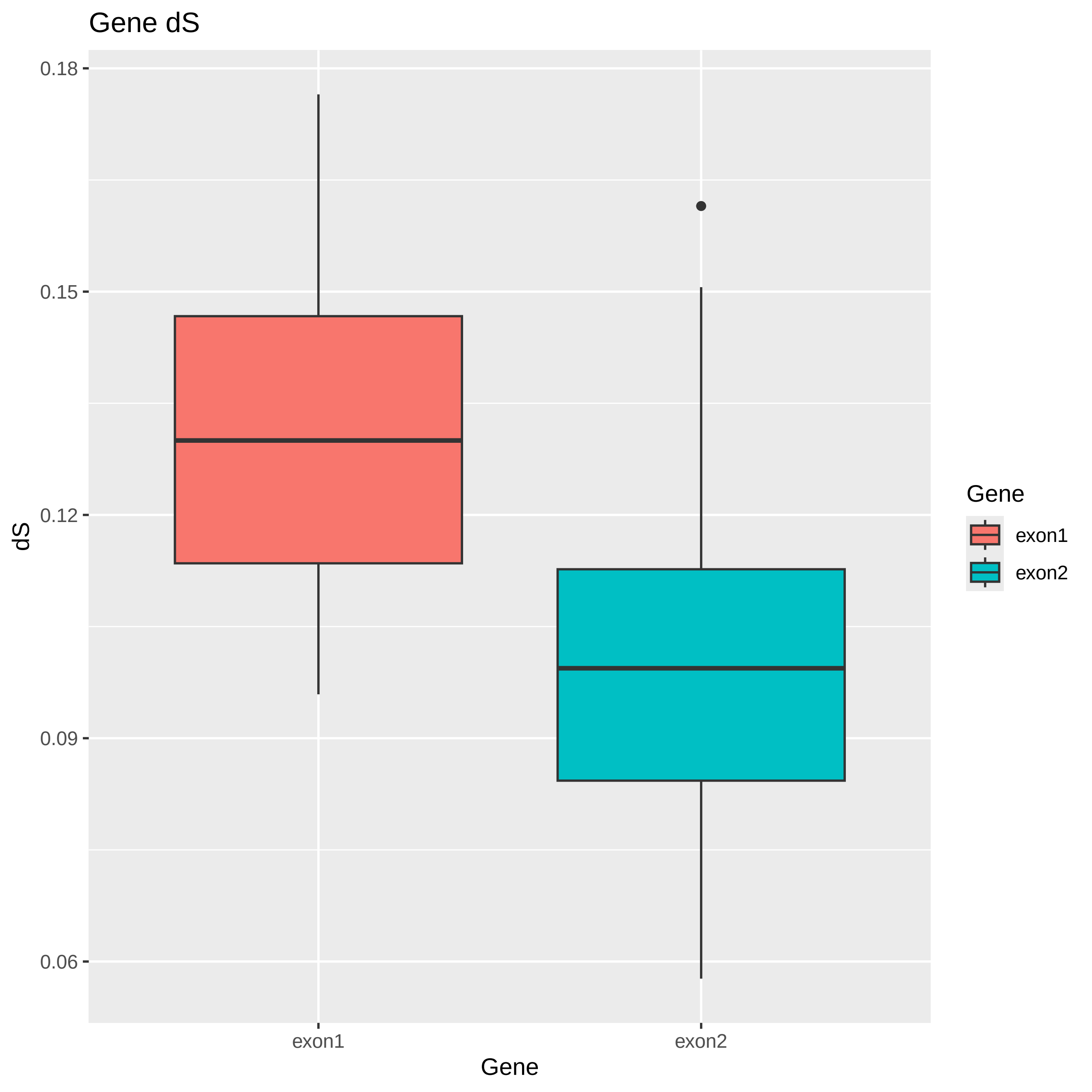

### exon1_dS_per_tribe.pdf

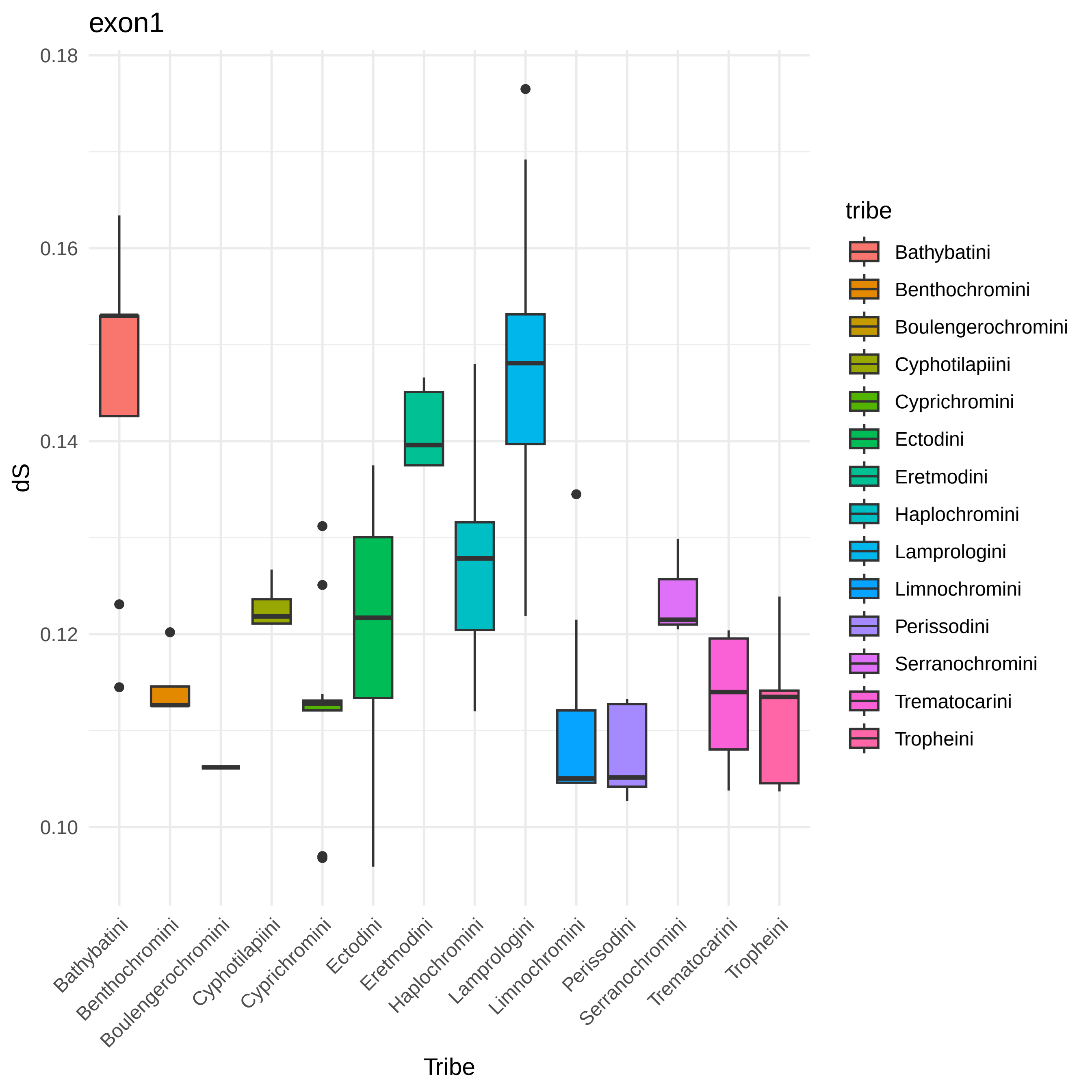

### exon2_dS_per_tribe.pdf

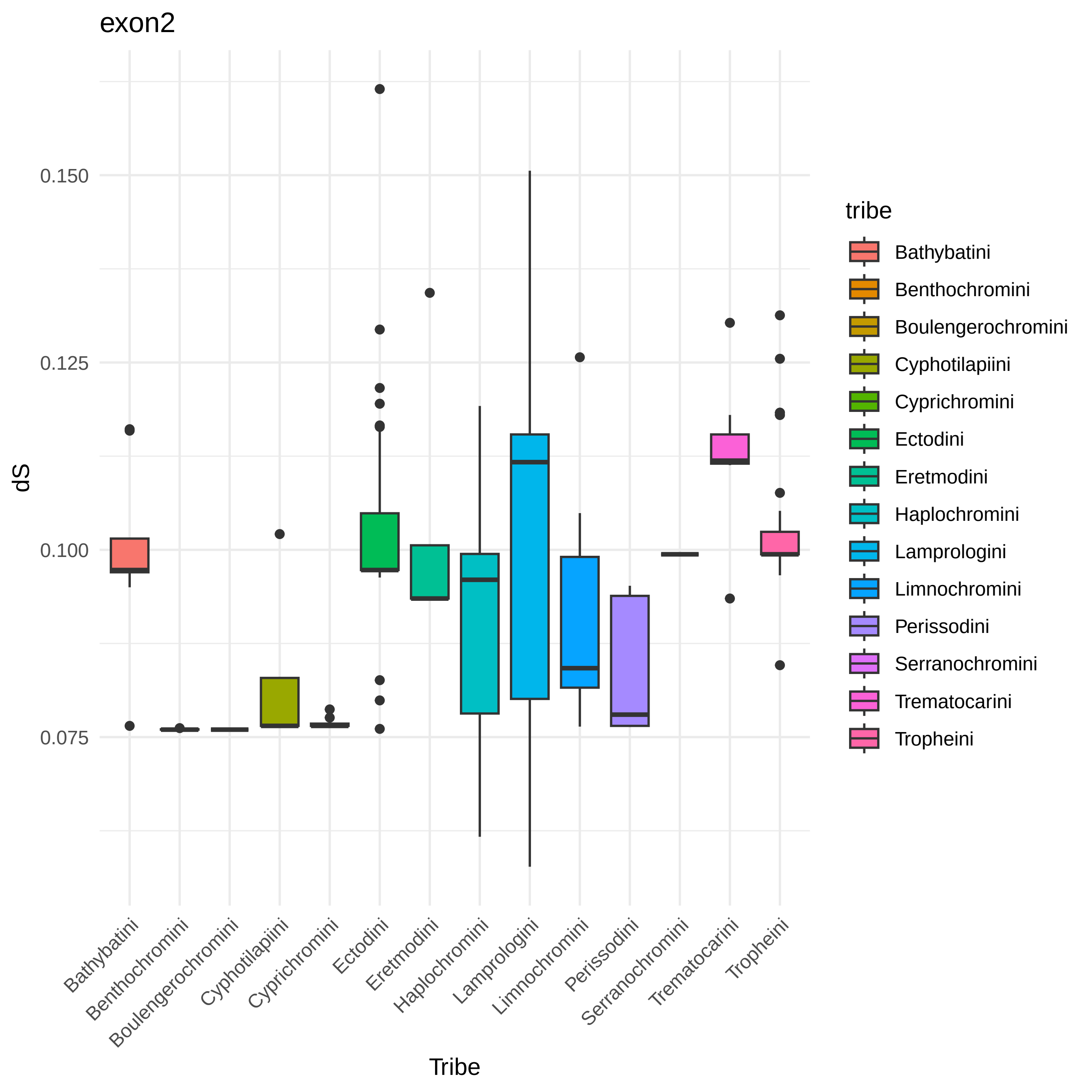

### OT_dS_per_tribe.pdf

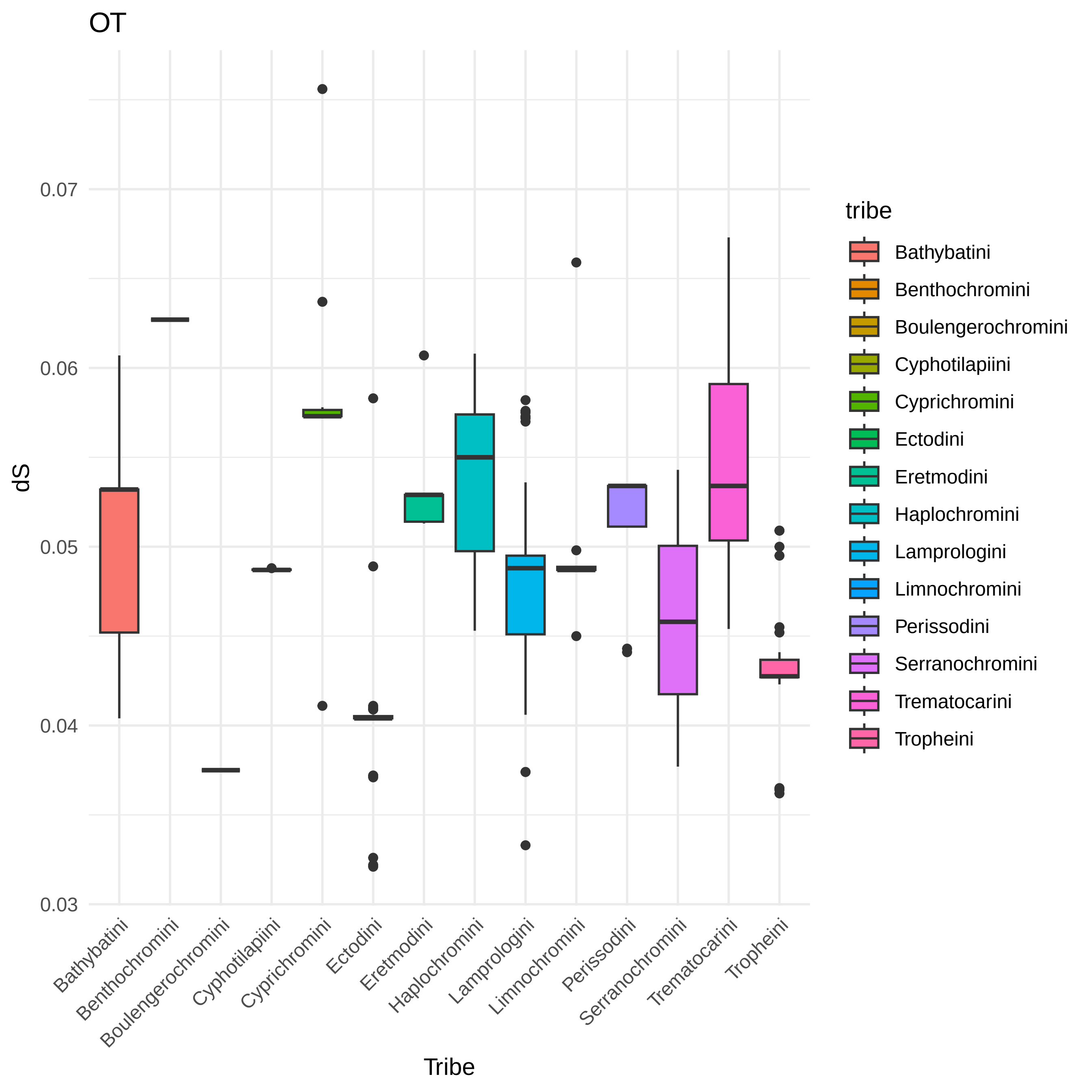

### OTRa_dS_per_tribe.pdf

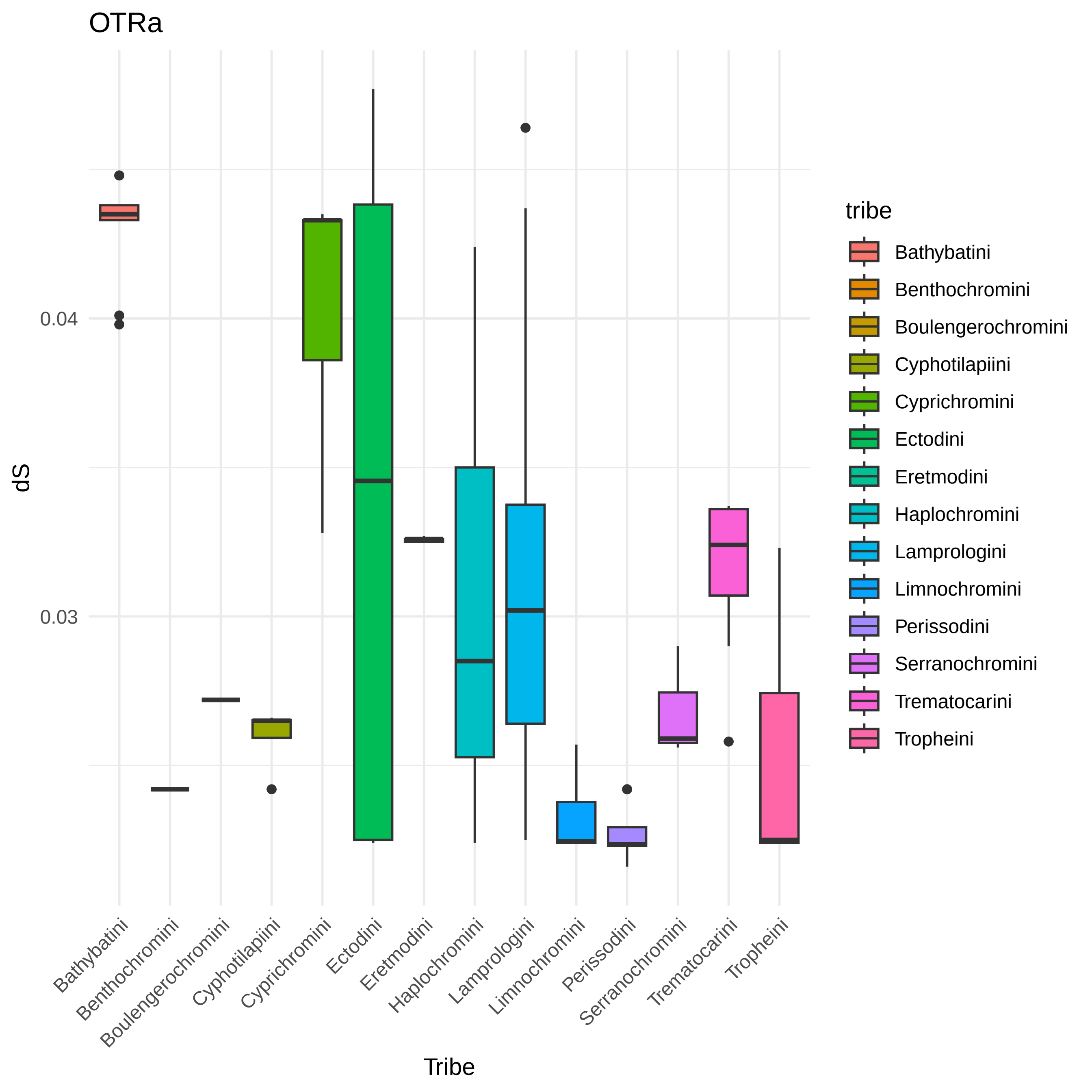

### OTRb.tr1_dS_per_tribe.pdf

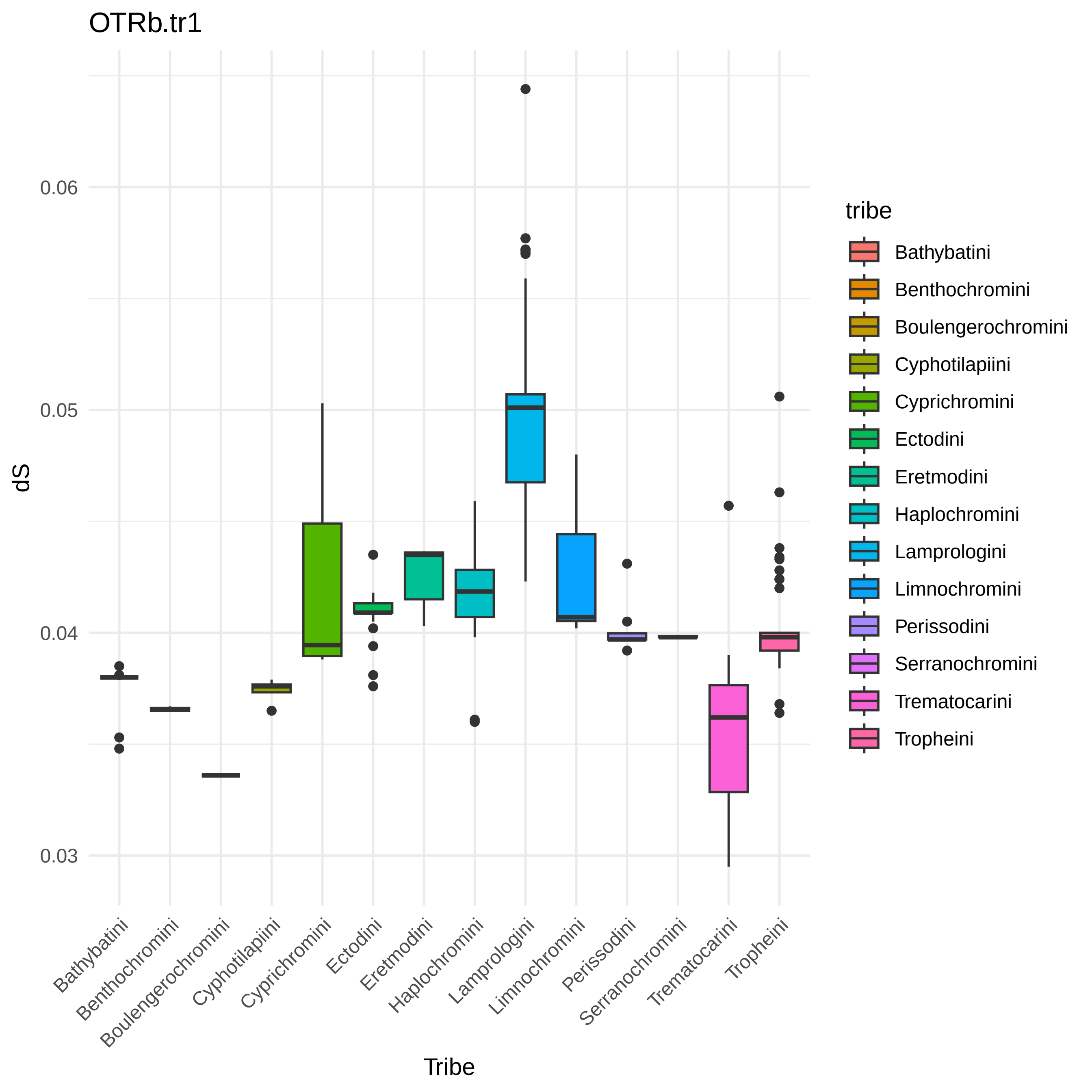

### OTRb.tr2_dS_per_tribe.pdf

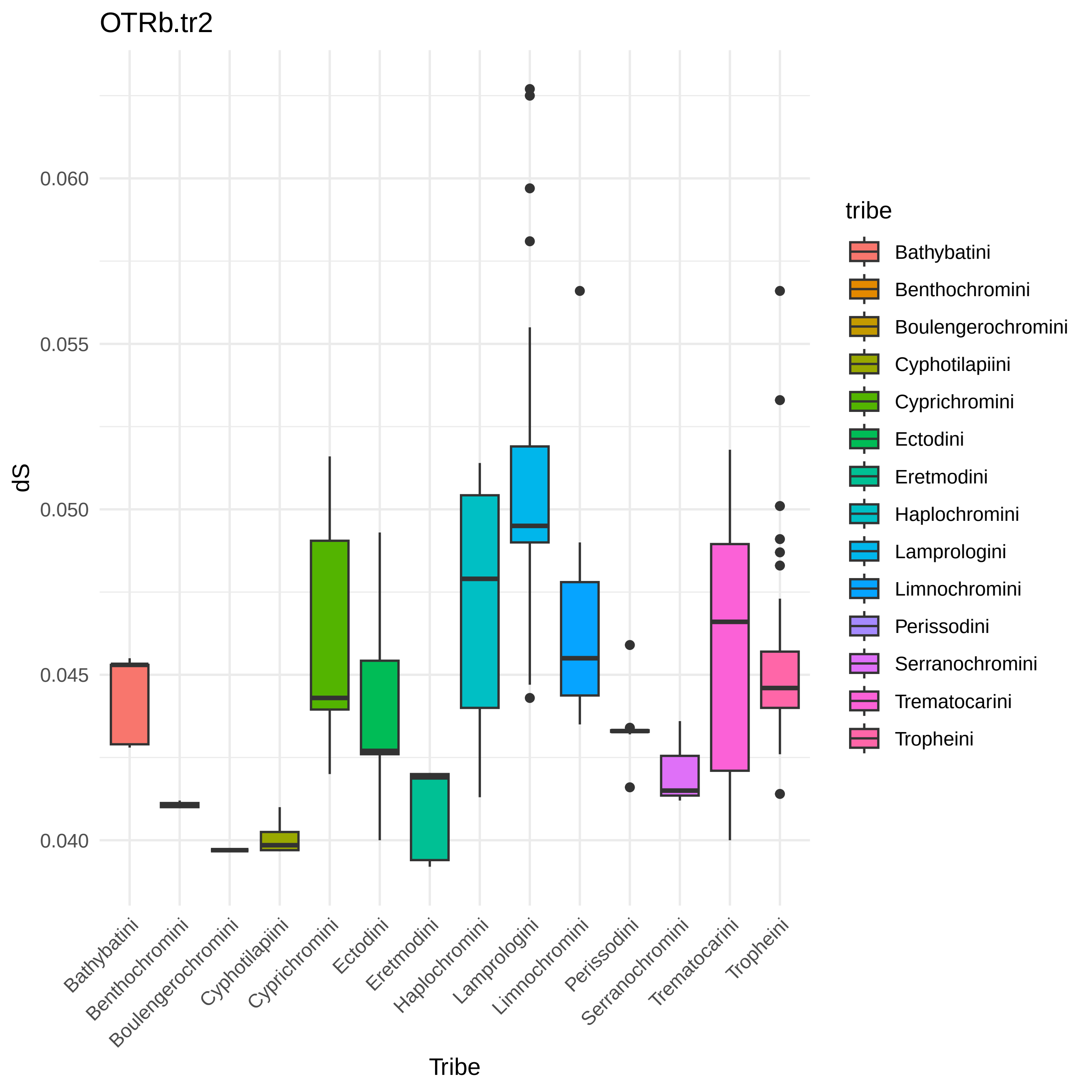

### OTRb.tr3_dS_per_tribe.pdf

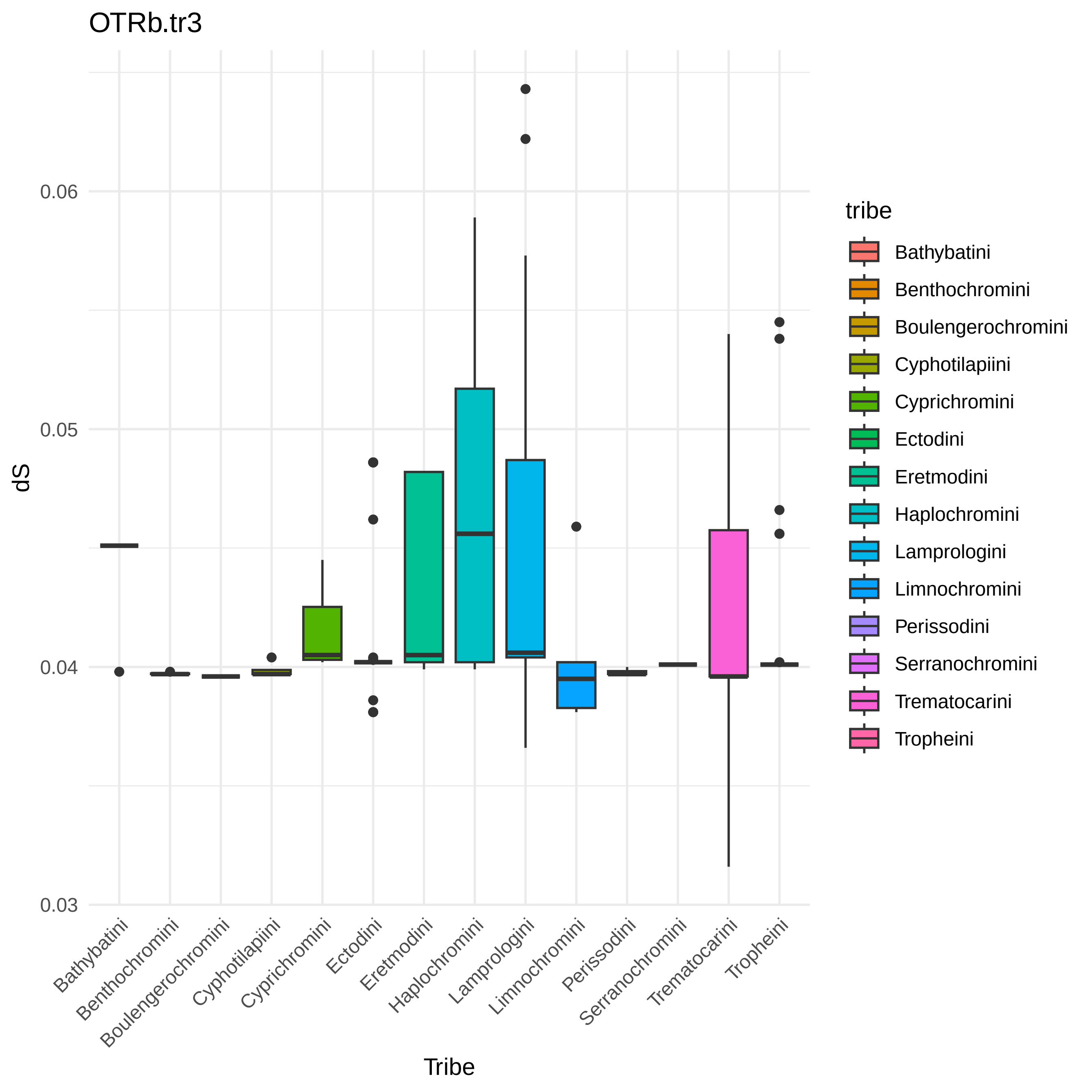

### Receptors Species Tree.pdf

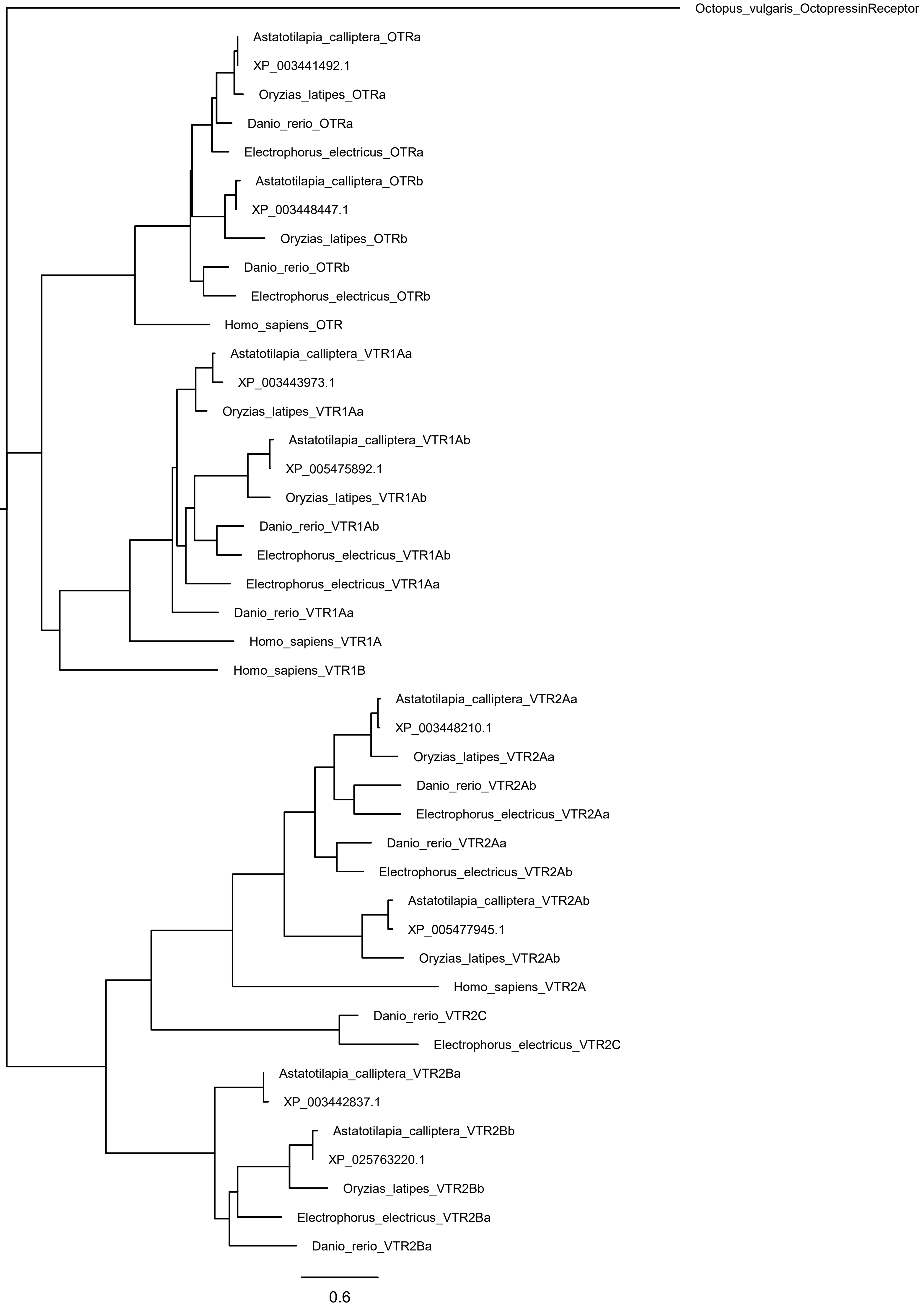

### VT_dS_per_tribe.pdf

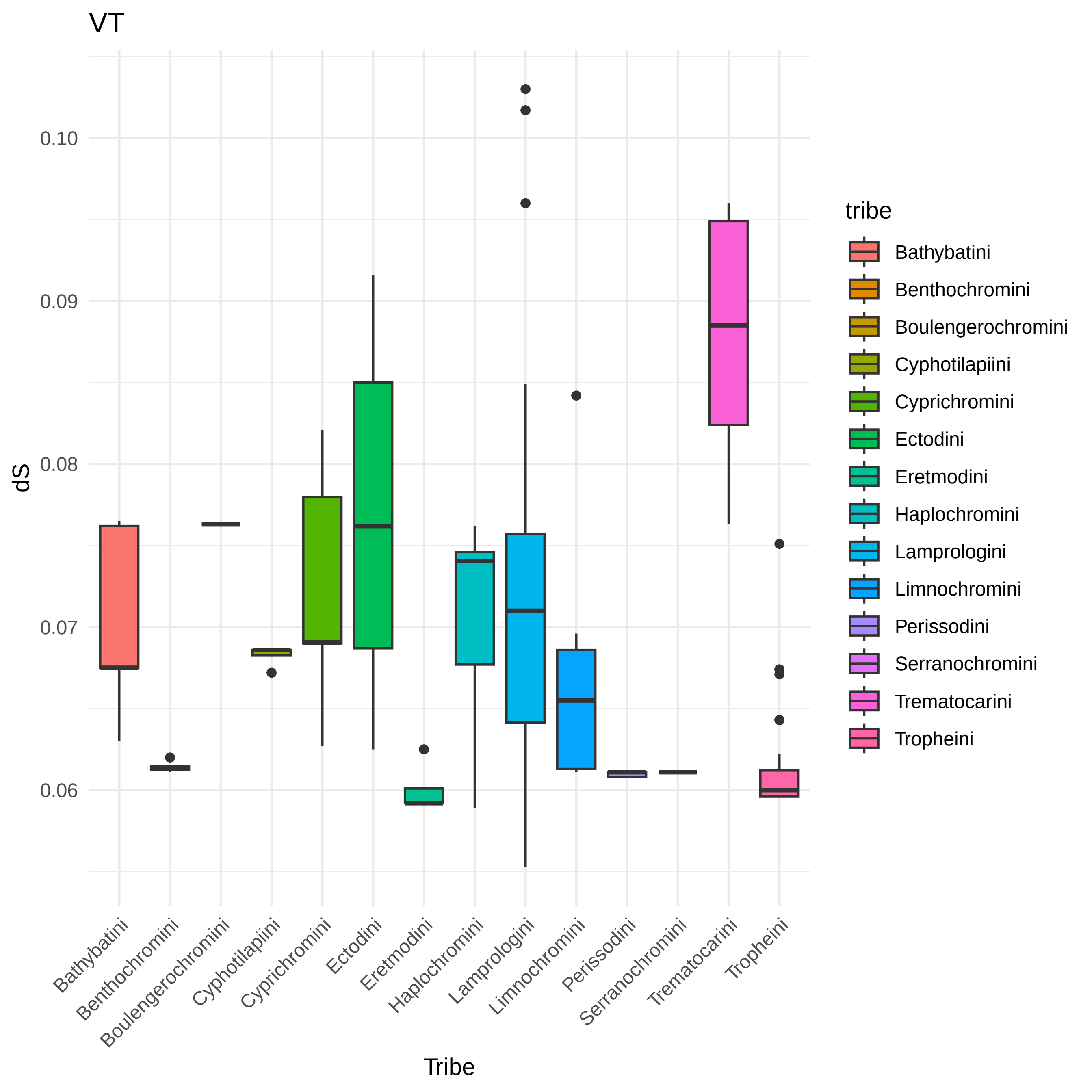

### VTR1Aa.tr1_dS_per_tribe.pdf

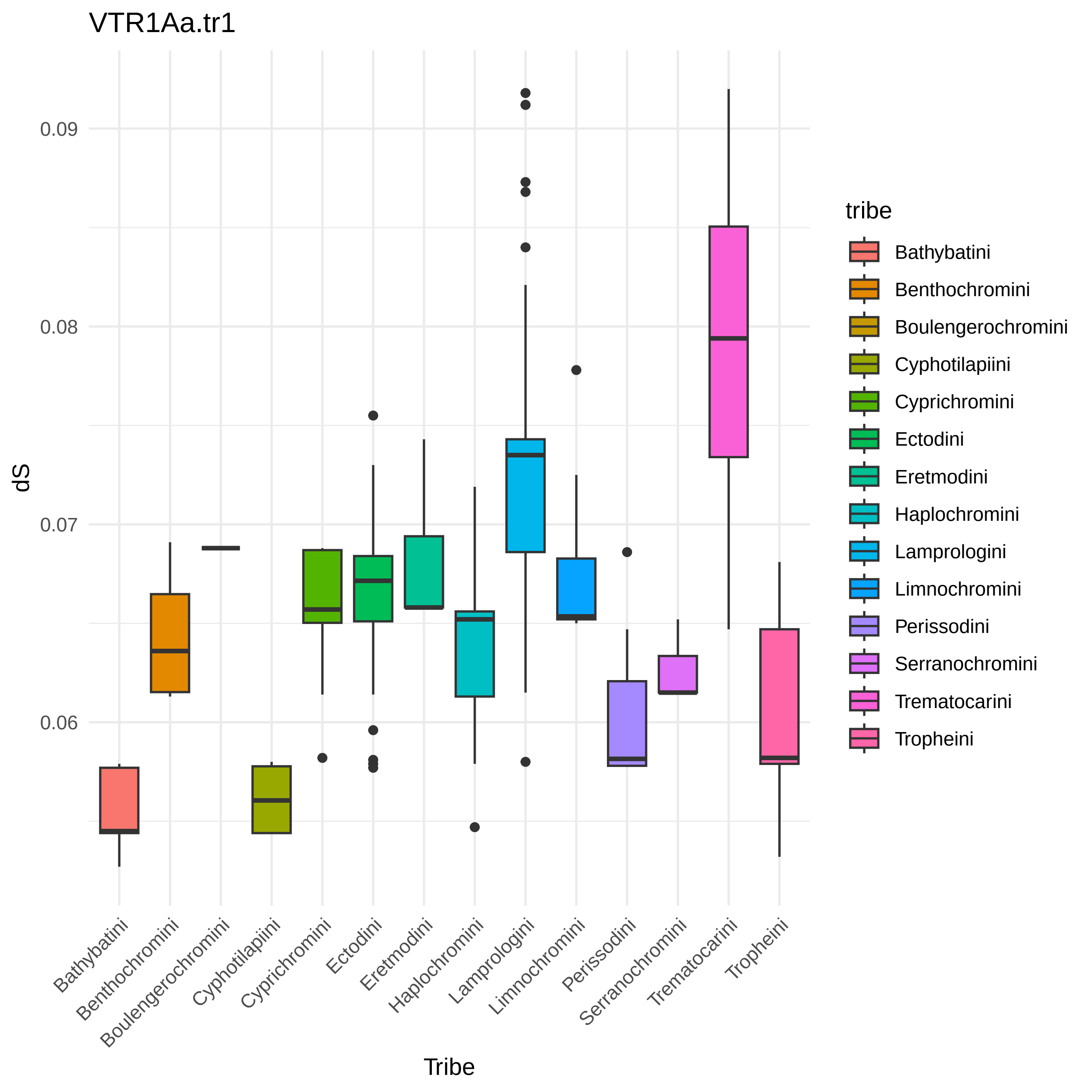

### VTR1Aa.tr2_dS_per_tribe.pdf

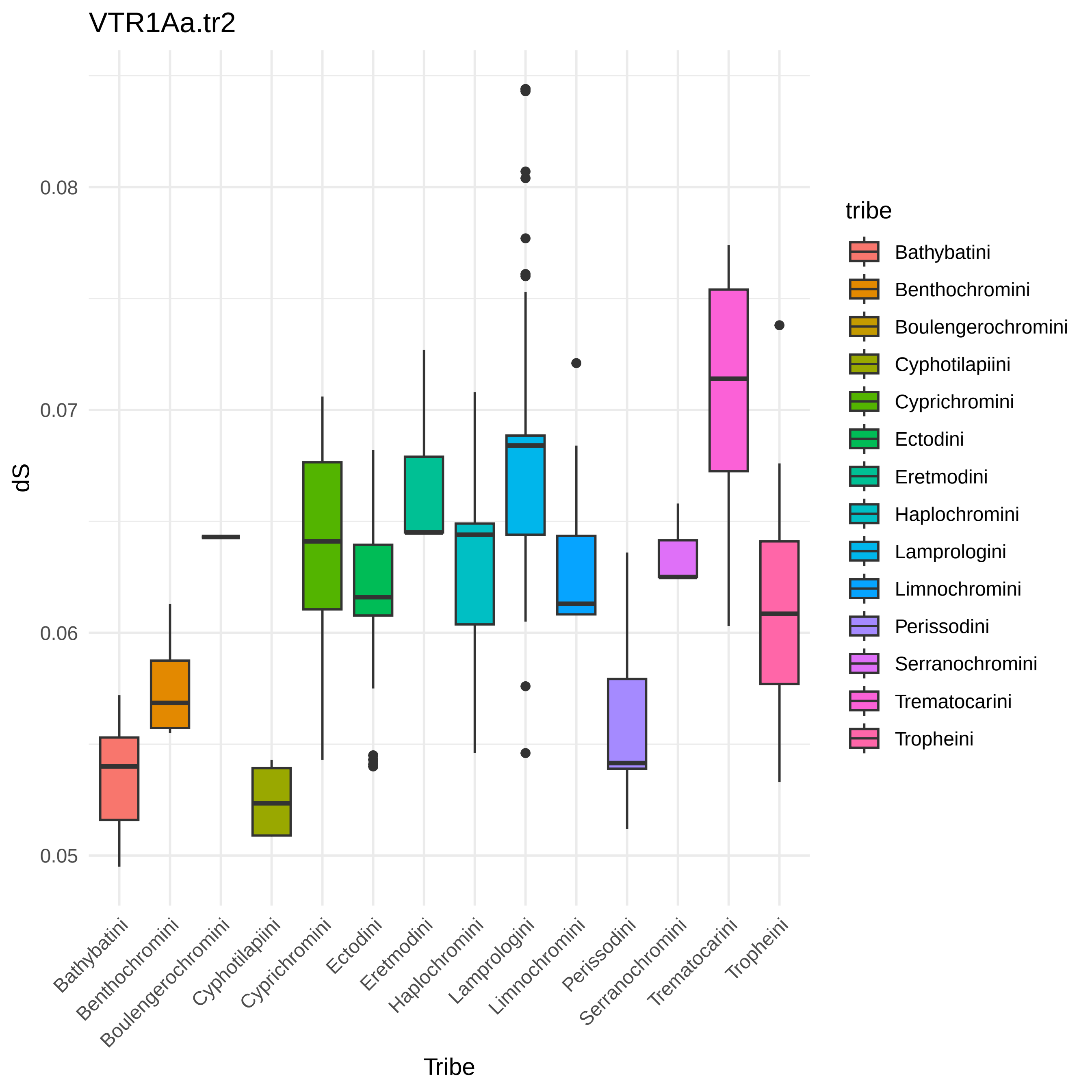

### VTR1Ab_dS_per_tribe.pdf

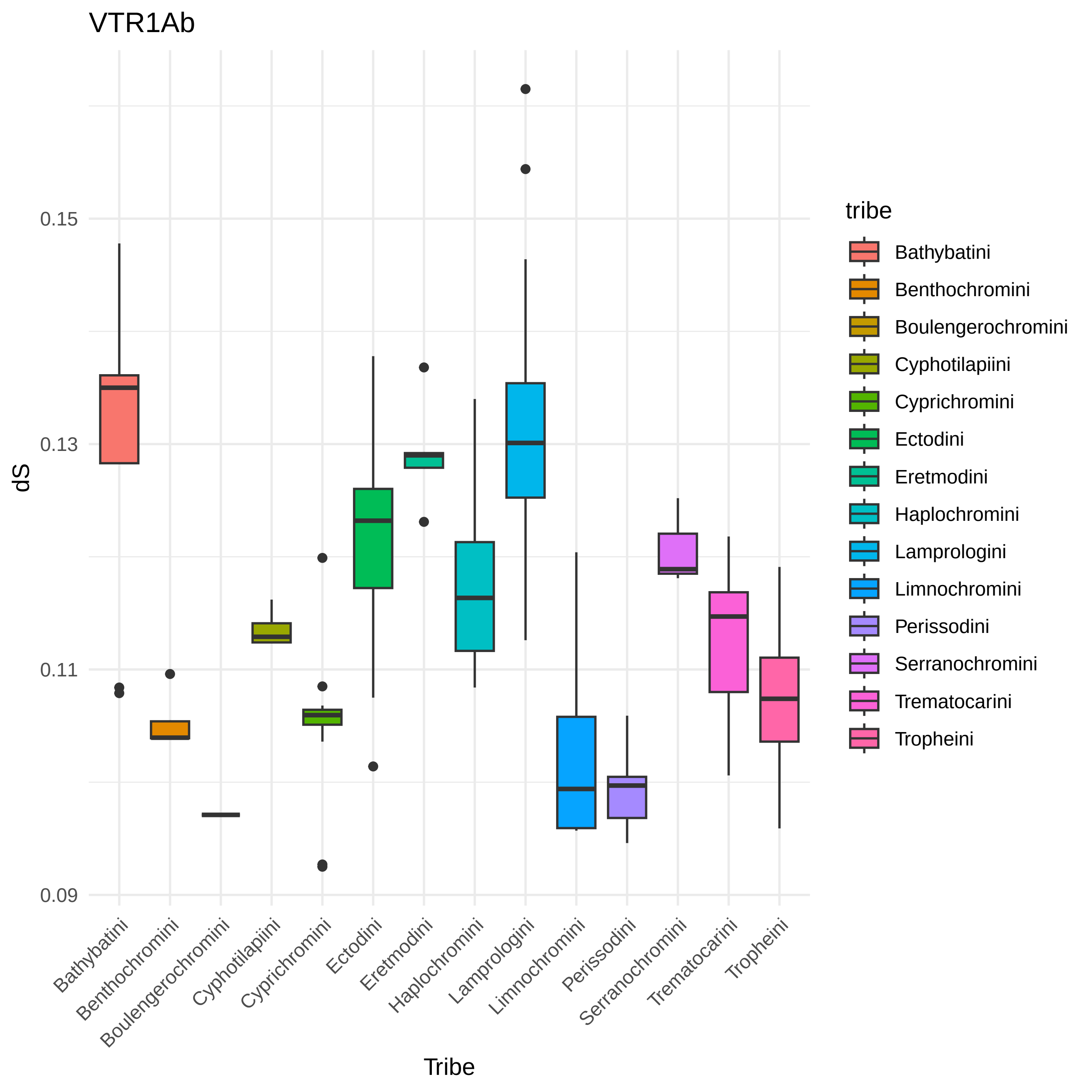

### VTR2Aa_dS_per_tribe.pdf

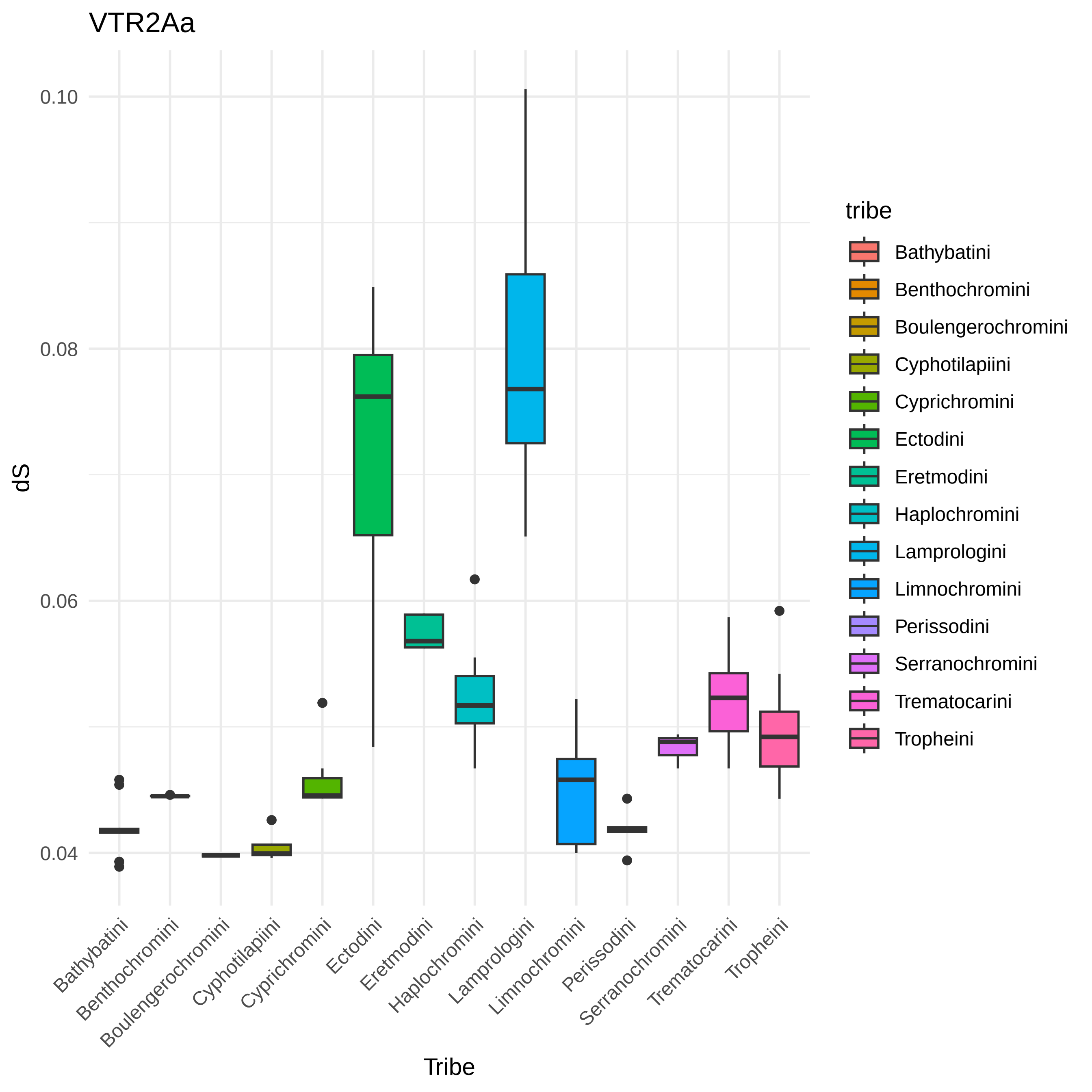

### VTR2Ab.tr1_dS_per_tribe.pdf

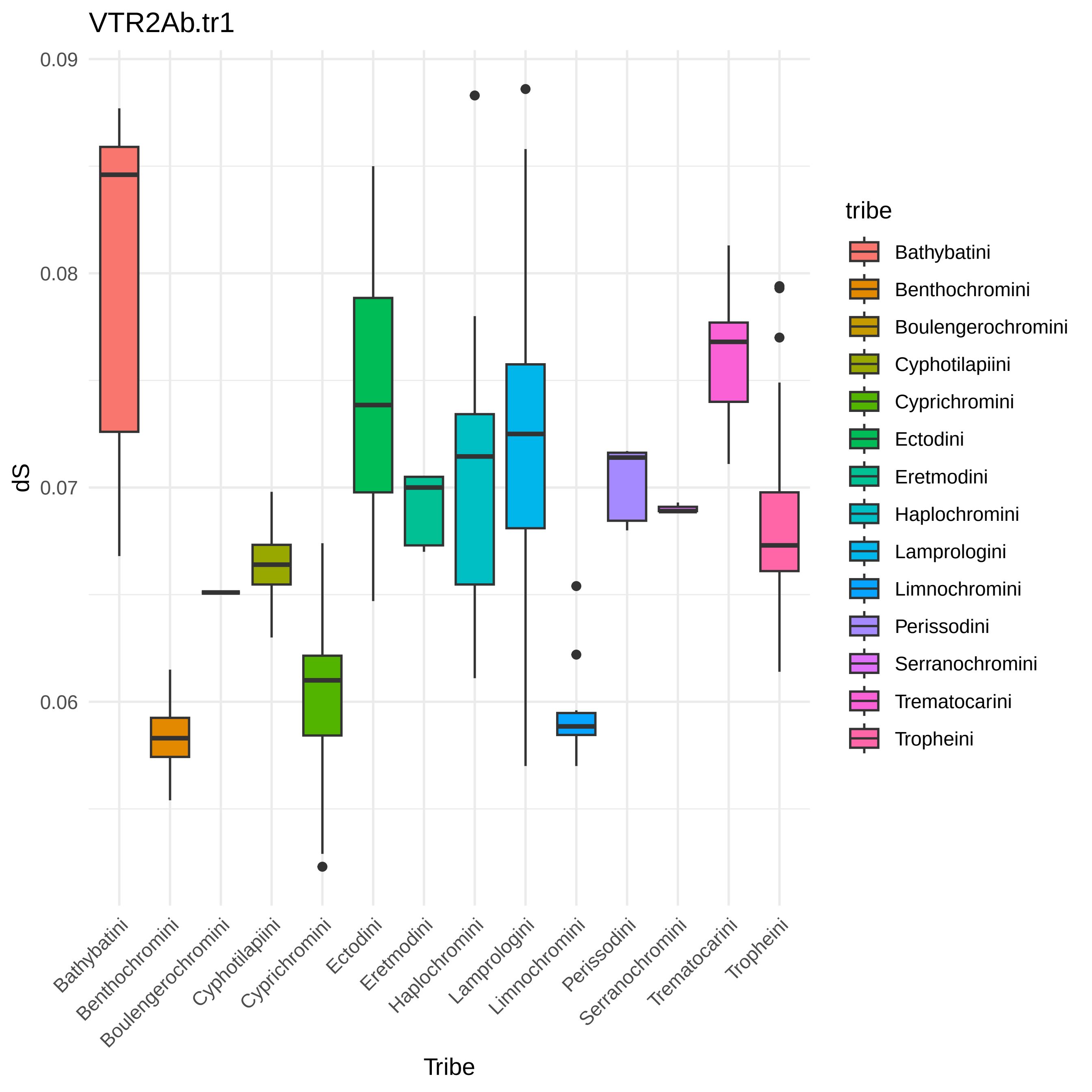

### VTR2Ab.tr2_dS_per_tribe.pdf

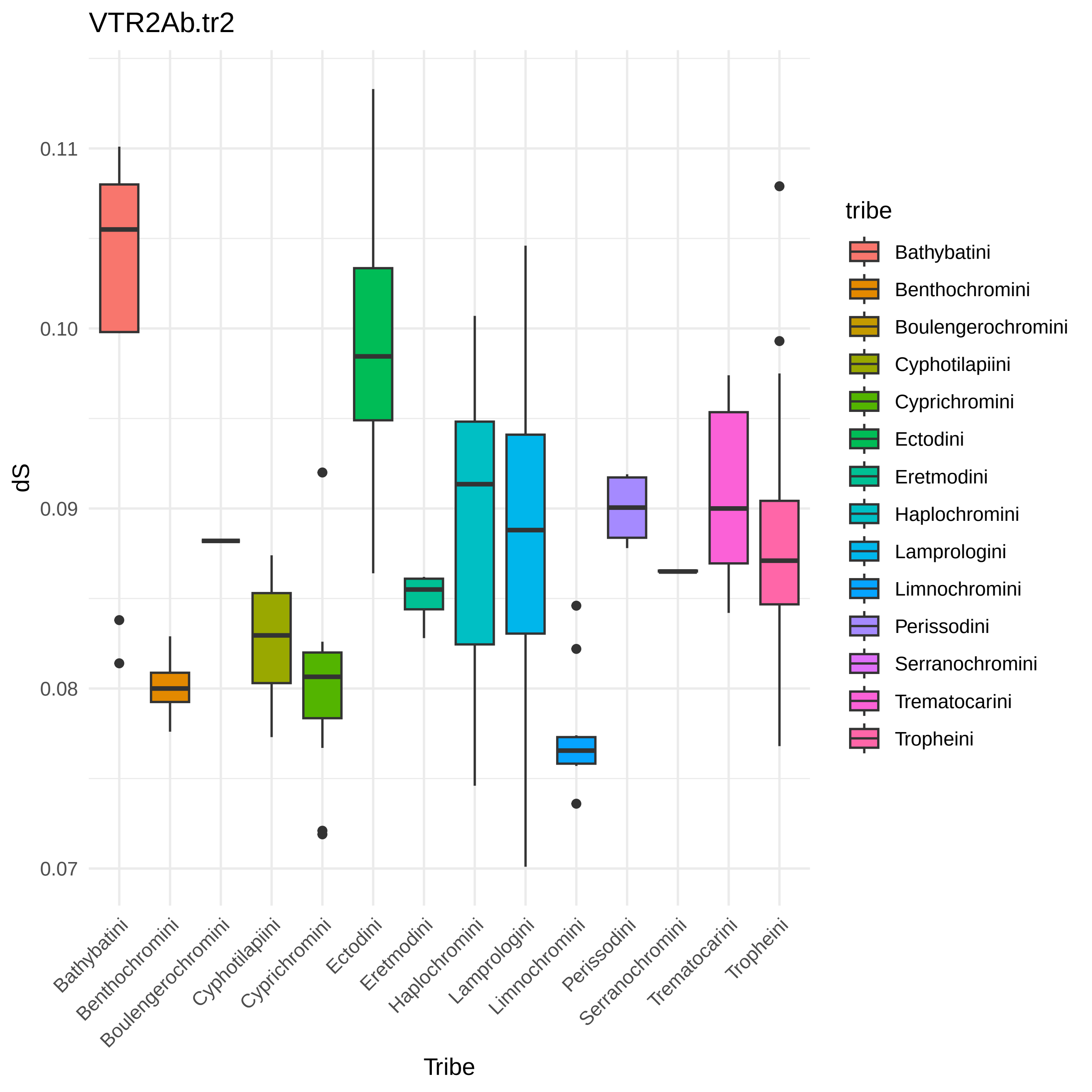

### VTR2Ab.tr3_dS_per_tribe.pdf

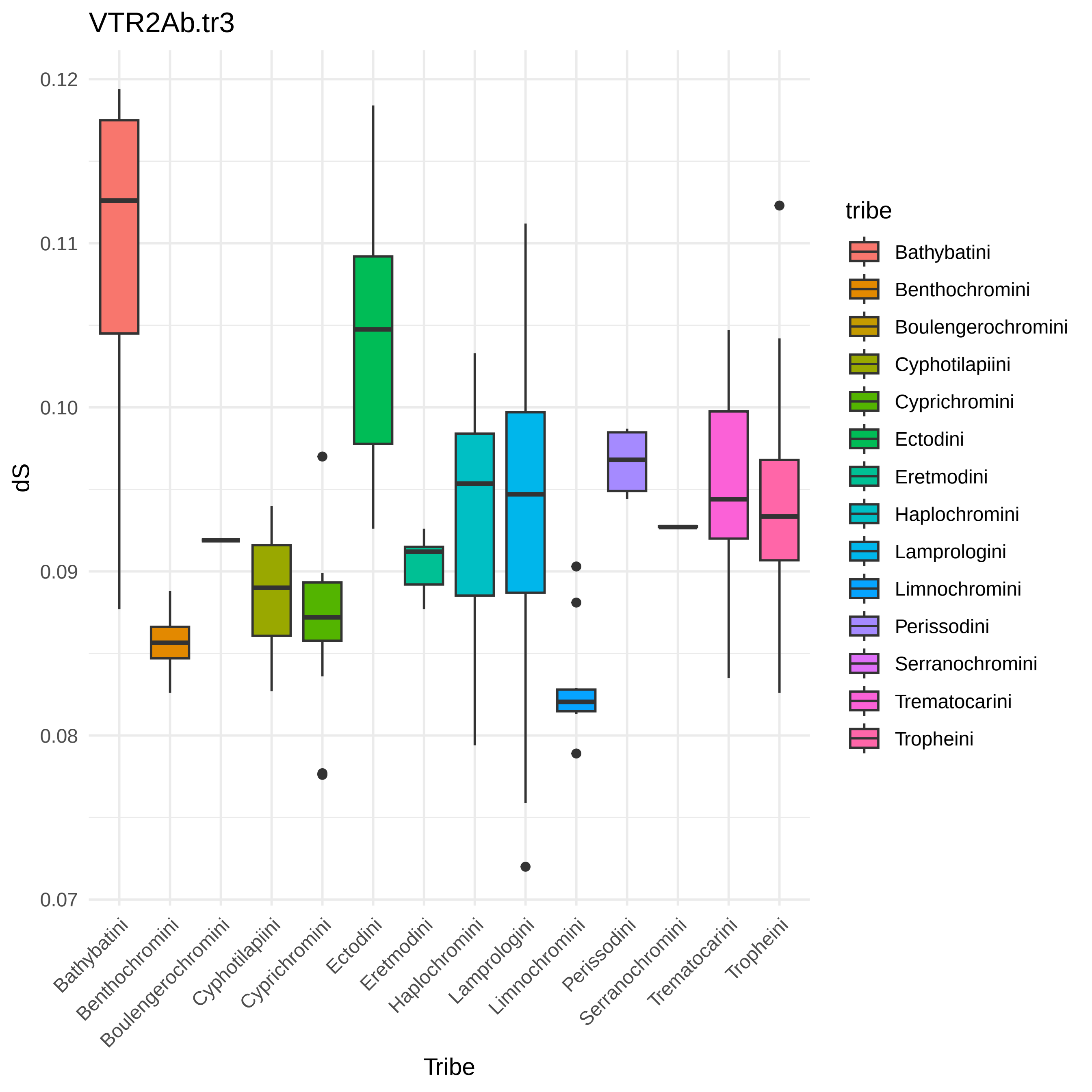

### VTR2Ba_dS_per_tribe.pdf

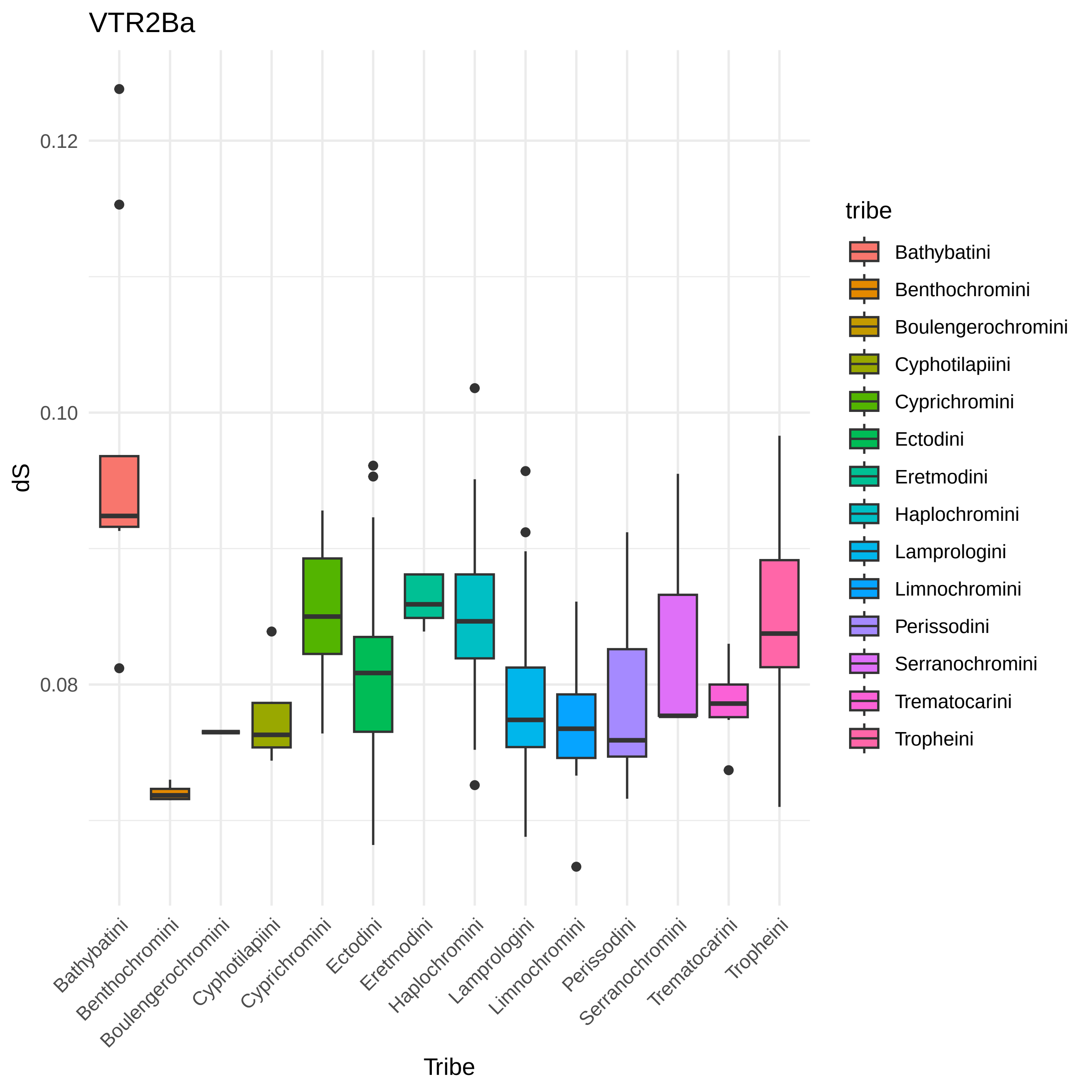

### VTR2Bb.tr1_dS_per_tribe.pdf

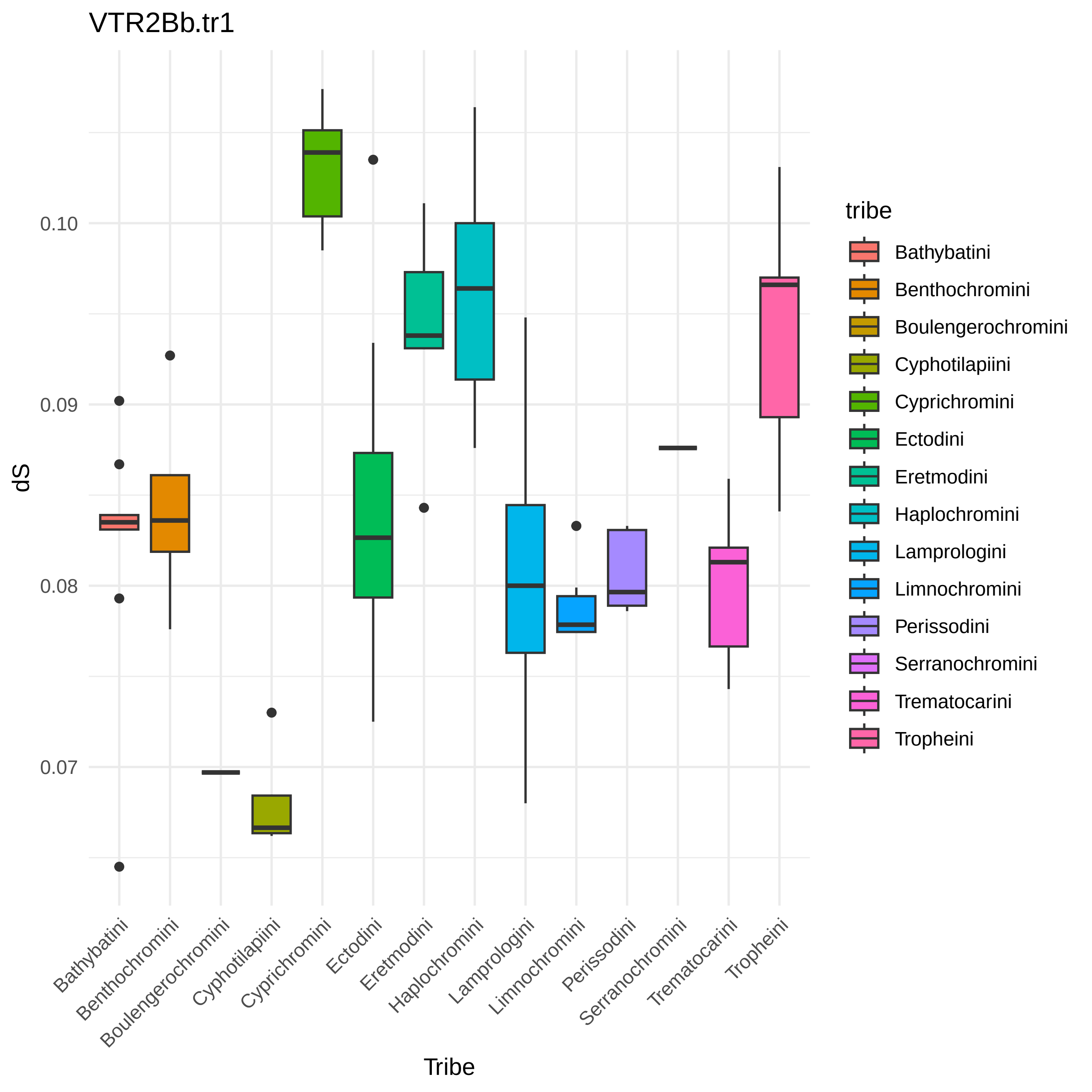

### VTR2Bb.tr2_dS_per_tribe.pdf

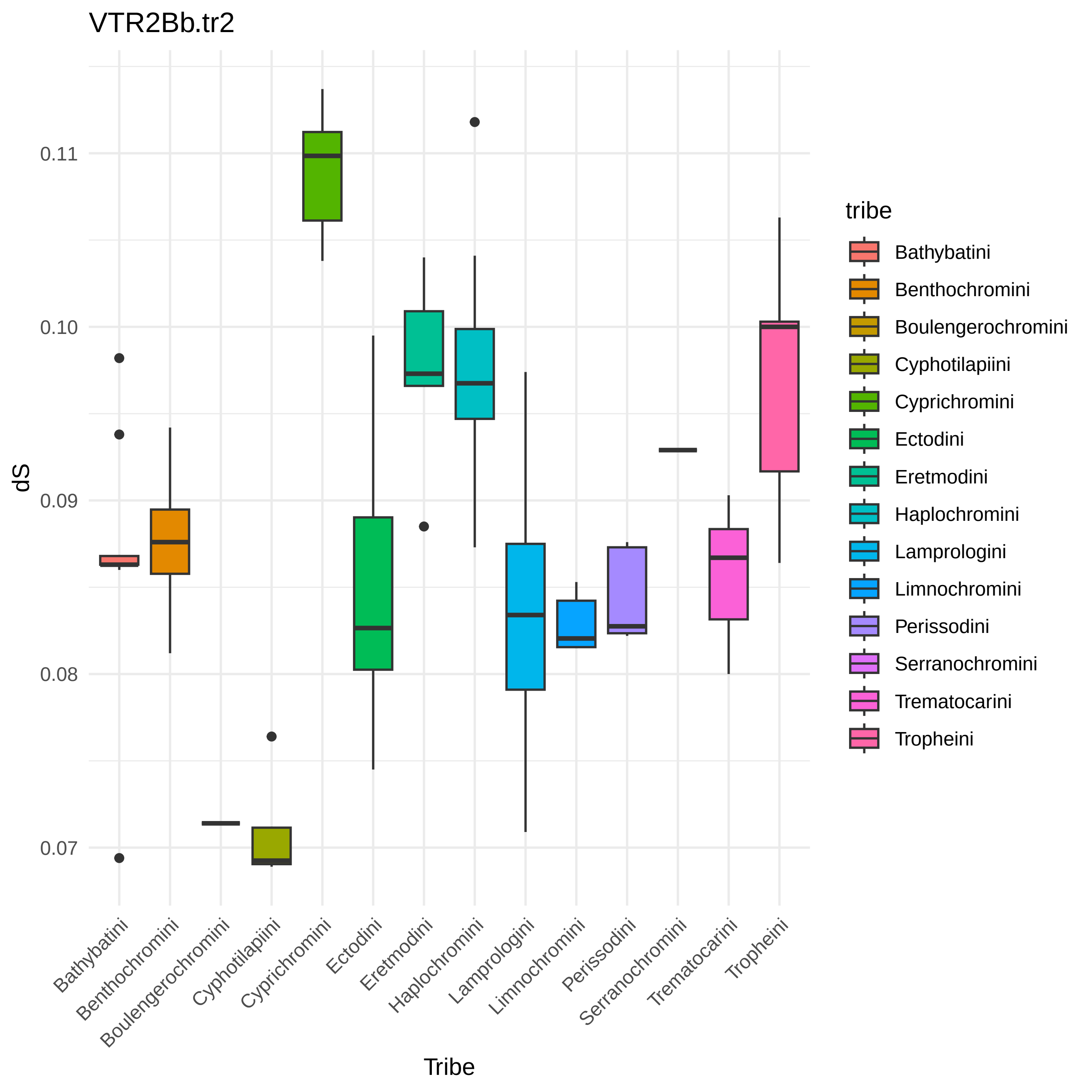
